# Human heart organoids reveal a regenerative strategy for mitochondrial disease

**DOI:** 10.64898/2025.12.18.695153

**Authors:** Minh Duc Pham, Arka Provo Das, Yue Wang, Despina Stefanoska, Ting Yuan, Chaonan Zhu, Meiqian Wu, Alfredo Cabrera-Orefice, Luka Nicin, Shane Bell, Mathias Langner, Ulrich Gärtner, Oliver Russell, Jonathan Ward, Mirko Peitzsch, Wesley Abplanalp, Maria Duda, Yu Hao, Phillip Grote, Marek Bartkuhn, Ilka Wittig, Peter Mirtschink, Andreas Zeiher, Stefanie Dimmeler, Jaya Krishnan

## Abstract

The human heart is among the most complex tissues to replicate in vitro, with vascular, neuronal, and immune elements shaping its development and function. Here we describe *cardiomorphs*, self-organising human cardiac organoids that recapitulate the cellular diversity, structural organisation, vascularisation, and innervation of the myocardium and mature along a developmental trajectory from early cardiogenesis to adult tissue. Using patient-derived *cardiomorphs*, we establish the first three-dimensional human tissue model of Kearns–Sayre syndrome (KSS), a rare mitochondrial disorder characterised by large-scale mtDNA deletions. KSS-cardiomorphs faithfully reproduce disease-associated metabolic, contractile, and ultrastructural hallmarks. Leveraging this platform, we identify Betaxolol, an FDA-approved selective β1-adrenergic antagonist, as a modulator of mitochondrial quality control. Betaxolol increases intracellular oxygenation, selectively eliminates dysfunctional mitochondria via mitophagy, and promotes biogenesis of functional organelles, restoring contractility in KSS tissues. This dual-action, mutation-agnostic mechanism suggests a therapeutic principle with broad relevance to mitochondrial disease, cardiac pathology, and age-associated decline.

## Introduction

Mitochondrial disorders are among the most devastating and least treatable conditions in medicine. They arise from mutations or deletions in the mitochondrial genome or in nuclear genes encoding mitochondrial proteins and affect an estimated 1 in 5000 individuals worldwide^1^. Despite their rarity, these disorders impose a profound clinical burden because mitochondria are central to energy metabolism, signalling, and cell survival. In tissues with high energetic demand, such as the heart, skeletal muscle, brain, and endocrine organs, mitochondrial dysfunction manifests as progressive multisystem dysfunction^2^.

Kearns–Sayre syndrome (KSS) epitomises these challenges. KSS is a rare mitochondrial disorder caused by large-scale deletions of mitochondrial DNA (mtDNA), typically spanning 5–9 kb and encompassing genes critical for oxidative phosphorylation (OXPHOS). Patients typically present in childhood or adolescence with progressive external ophthalmoplegia, pigmentary retinopathy, muscle weakness, and neurological deficits^7,8^. Critically, the heart is among the most affected organs with up to 84% of patients developing cardiac complications including conduction block, arrhythmias, and hypertrophic cardiomyopathy, with 23% dying from sudden cardiac death^7,8^. KSS is characterized by increasing disease progression with age, concomitant with increasing mutant heteroplasmy (i.e. the proportion of mutant to wild-type mitochondrial genomes) driving increasing severe phenotypes^2^. Current therapies are largely symptomatic. Whilst implantation of a pacemakers can prevent arrhythmic death in patients, and supplements such as coenzyme Q10 can be beneficial, neither approach addresses the root cause of KSS and the compounding impact of dysfunctional mitochondria accumulation with age^3^.

The lack of appropriate human models has hindered therapeutic development. Patient biopsies are exceedingly scarce, especially from the heart, and in vitro models have traditionally relied on isolated fibroblasts or reprogrammed cardiomyocytes. Although useful, these systems fail to recapitulate the architectural, metabolic, and multicellular complexity of human cardiac tissue. Over the past decade, tissue engineering and organoid technologies have sought to overcome this gap. However, current cardiac organoids or spheroids suffer significant limitations. Most cardiac models generate cardiomyocytes alone, or mixtures of cardiomyocytes with fibroblasts and occasionally endothelial cells^4–13^. Even with endothelial inclusion, these tissues seldom form perfusable vascular networks with proper mural cell coverage, and they rarely integrate neuronal or immune components^14–16^. This lack of minority lineages is not trivial: as neurons regulate contractility and rhythm, macrophages contribute to remodelling and repair, and stromal interactions orchestrate paracrine signalling^17–20^. Without these cell fractions, organoids show restricted growth, impaired functional maturation, and blunted stress responses, which in turn compromise their value for disease modelling and drug testing^17–20^. The absence of minority cell types may explain why many cardiovascular drug candidates fail in clinical trials, as preclinical models do not capture the multicellular physiology of the heart, leading to false positives in vitro that collapse under the complexity of human biology^21^. This results in a high attrition rate for cardiovascular therapies, with billions invested in compounds that never achieve clinical benefit.

To overcome these barriers, we developed cardiomorphs (CDMs): self-organising, induced pluripotent stem cell (iPSC)-derived human cardiac organoids that incorporate the full cellular diversity of the myocardium. CDMs contain not only cardiomyocytes but also endothelial cells, mural cells, neurons, macrophages, and stromal elements. These lineages organise into vascularised, innervated, immune-competent tissues that progress through a developmental trajectory from early cardiogenesis to adult-like myocardium. Benchmarking against mouse and human transcriptomes confirms stage-matched maturation.

CDMs are dynamic and undergo morphological and metabolic transitions, shifting from glycolysis to oxidative metabolism, and display adult-like contractility, drug responses, and capacity for vascular integration with native heart tissue, thus serving as a faithful human surrogate for cardiac physiology and pathology. Utilising this platform, we modelled Kearns-Sayre syndrome for the first time in three-dimensional human cardiac tissue. Using patient-derived iPSCs harbouring canonical mtDNA deletions, we generated KSS-CDMs that faithfully reproduced hallmark features: impaired OXPHOS, reduced ATP production, mitochondrial disorganisation, pathological hypertrophy, lipid accumulation, neurogenic deficits, and abnormal contractility. Mechanistic studies revealed destabilisation of MICOS–SAMM supercomplexes, essential for cristae architecture, and a loss of mitochondrial “microregeneration” (the capacity of postmitotic cells to repair and replace OXPHOS subunits over time). These findings establish CDMs not only as models of mitochondrial disease but also as discovery engines for pathophysiological insight. Further, CDMs provided a platform for therapeutic discovery. Recognising that KSS involves progressive expansion of mutant mitochondria in vivo but selective loss of mutants in hyperoxic culture, we hypothesised that cellular oxygenation states influence mitochondrial selection. We therefore performed a targeted pharmacological screen of FDA/EMA-approved drugs known to alter oxygen metabolism, including β-adrenergic antagonists and agonists, antiarrhythmics, and antidepressants. While many compounds proved toxic or ineffective, the β-adrenergic antagonists Betaxolol (β1-selective) and Nadolol (non-selective) emerged as potential potent therapeutics for KSS-CDM viability and function. Mechanistic assays revealed that Betaxolol and Nadolol suppressed oxygen consumption, elevated intracellular oxygenation, and triggered a dual program of selective mitophagy of depolarised and dysfunctional mitochondria and the induction of biogenesis of polarised, functional mitochondria. This mitophagy-biogenesis coupling restored mitochondrial polarisation, reduced mutant mtDNA load, and abrogated the cardiac contractile dysfunction associated with KSS.

Importantly, this mutation-agnostic strategy does not rely on correcting the underlying mitochondrial or nuclear genetic lesion. Rather, it exploits competitive fitness among mitochondrial populations, creating an intracellular environment in which healthy mitochondria thrive and mutants are selectively eliminated. The conceptual significance of this mechanism could be profound, as it suggests a generalisable regenerative principle that could apply across diverse diseases characterised by mitochondrial dysfunction, from rare syndromes like KSS to common disorders of heart disease, neurodegeneration, metabolic syndrome, and even physiological ageing. Thus, the development of cardiomorphs and their application to KSS provide not only a new experimental platform but also a new therapeutic paradigm wherein it may be possible to restore function in diseases long considered untreatable simply by reshaping intracellular oxygenation landscapes.

## Results

### Derivation and developmental benchmarking of cardiomorphs

We aimed to establish conditions to generate human cardiac organoids that capture both the multicellular architecture and the developmental progression of the myocardium. Thus, induced pluripotent stem cells (iPSCs) were directed to the mesoderm lineage under through staged signalling conditions that combined Activin A and BMP4 with FGF2 and controlled Wnt/β-catenin modulation, and ascorbic acid 2-phosphate was included to promote cardiogenic competence during lineage transition (**Fig. 1A; Extended Data Fig. 1A-F; Extended Data Fig. 2A**)^22–26^. At day 0, cells aggregated in low-adhesion microwells and formed uniform spheroids that initiated self-organization. Early aggregates exhibited epithelialised surfaces and compact cores, consistent with mesodermal specification, and developed focal contractions detectable by day 7 (**Fig. 1B; Extended Data Fig. 2B**)^27^. Coordinated contractions spanning the entire organoid were observed by day 10 and increased in amplitude through day 35 (**Extended Data Fig. 2B; Extended Data Video 1**).

**Figure 1.**
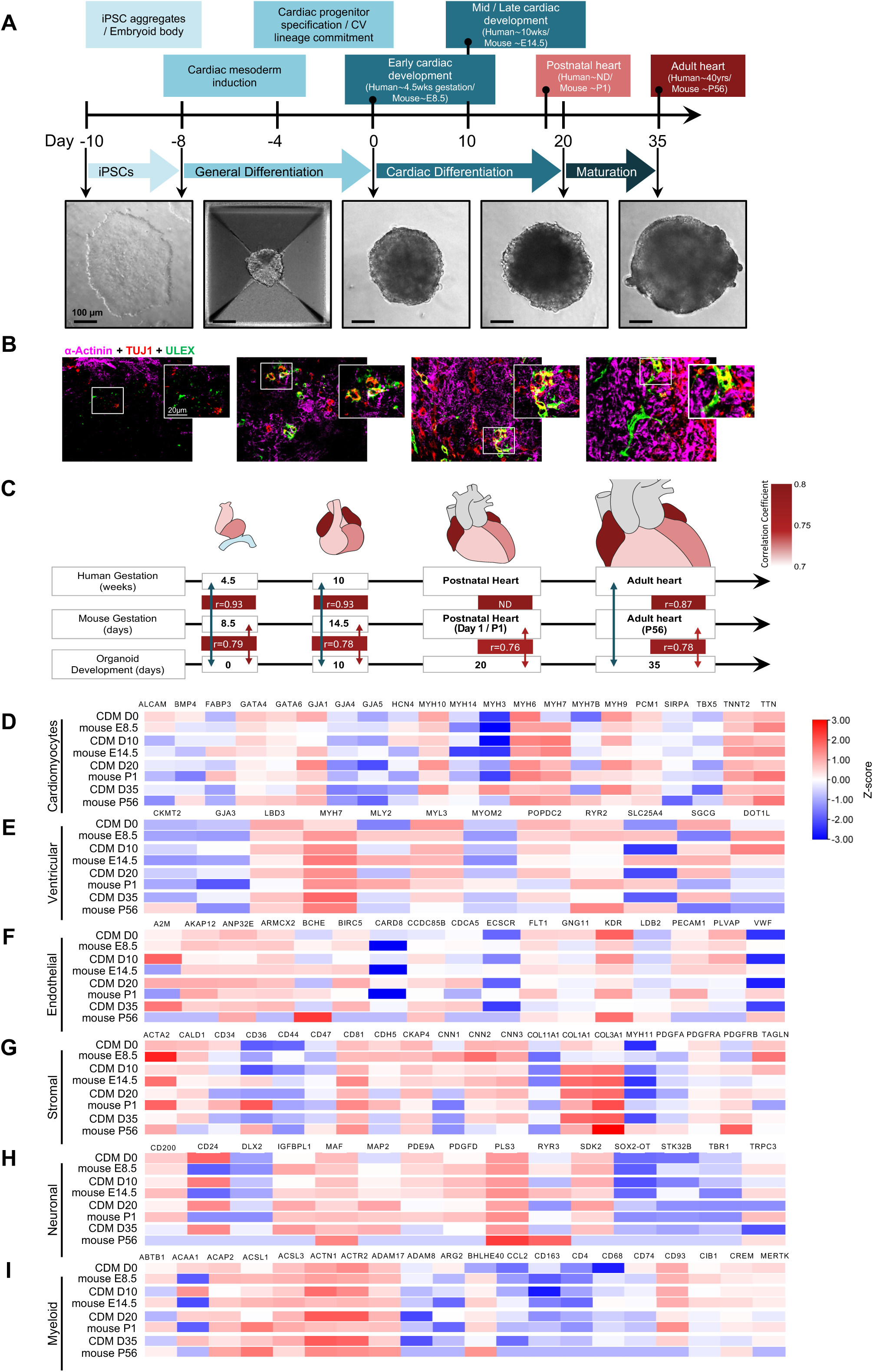
Cardiomorphs recapitulate human cardiac development. **A,** Schematic representation of the differentiation timeline with corresponding brightfield images of CDM growth, as benchmarked to equivalent stages of mouse heart development. Scale bar, 100 µm. **B,** Representative images of CDM at different development stages as corresponding to schematics in (**A**). CDM were stained with nuclear marker (DAPI − blue), endothelial marker (ULEX − green), neuronal marker (TUJ1 − red) and cardiac marker (α-Actinin − magenta). Scale bar, 40 µm; inlet scale bar, 20 µm. **C,** Pearson’s correlation between heart development gene expression pattern of CDM, and mouse and human heart development at corresponding stages. (*n* = 2 − 4 replicates per sample). **D-I,** Longitudinal CDM expression profiles of representative gene markers for (**D**) cardiomyocytes (CM), (**E**) ventricular lineage cells, (**F**) endothelial cells (EC), (**G**) stromal cells, (**H**) neuronal cells, and (**I**) myeloid cells at different development stages as corresponding to schematics in (**A**). (*n* = 2 replicates with 100 CDMs pooled per replicate; expression values are shown as z-scores for each gene).

Temporal transcriptome profiling revealed that cardiomorphs (CDMs) advance through discrete developmental stages aligned with human ontogeny. Bulk RNA-seq principal component analysis (PCA) and Pearson correlation benchmarking against published mouse and human cardiac reference sets indicated that day 10 organoids map to an early foetal state, whereas day 35 organoids align most closely with postnatal myocardium (**Fig. 1C**)^28,29^. Canonical stage markers followed expected trajectories: mesodermal regulators (e.g., MESP1, ISL1) peaked early and declined as sarcomeric genes (e.g., TNNT2, MYH6/MYH7, PLN) rose, together with Ca2+ handling genes (e.g., RYR2) (**Fig. 1D,E**). Cardiomyocytes exhibited a foetal-to-adult myosin heavy chain switch, maturation of ion-handling transcripts, and enrichment of structural modules consistent with sarcomere alignment (**Fig. 1D,E**). Endothelial clusters expressed PECAM1 and CDH5; mural clusters expressed ACTA2 and PDGFRB; neuronal clusters expressed TUBB3 (TUJ1) and MAP2; macrophages expressed CD68 and MRC1; fibroblast/stromal clusters expressed ECM genes (e.g., COL1A1, DCN) (**Fig. 1F-I**). Single-cell RNA-seq at day 35 confirmed its multicellular composition. Clustering resolved cardiomyocytes, endothelial cells, mural cells, neurons, macrophages, and stromal/fibroblast populations (**Extended Data Fig. 3A**). These proportions matched reported distributions in native human myocardium and remained stable across experiments (**Extended Data Fig. 3B**)^30^. Cell-type composition analyses highlighted largely consistent and parallel patterns of gene expression of the respective cell-types between CDMs and human ventricular biopsies (**Extended Data Fig. 3C,D**).

Immunofluorescence confirmed lineage identity and spatial organisation. Cardiomyocyte-specific α-actinin staining revealed striated myofibrils with apparent Z-line registration by day 35 (**Fig. 2A,B**). Endothelial cells formed VE-cadherin–positive tubes with consistent lumenal profiles, with mural coverage by NG2+ and SMA+ cells indicative of vessel maturation and stabilisation (**Fig. 2C,D**)^31–34^. TUJ1+ neurites extended from peripheral foci into the contractile tissue whilst CD68+ macrophages were distributed throughout, including near vascular structures (**Fig. 2E,F**)^31,35–37^. Three-dimensional confocal reconstruction showed that endothelial networks traverse the organoid and form long, directional segments, which is consistent with vessel guidance during morphogenesis (**Extended Data Fig. 4A,B**)^32–34^. Transmission electron microscopy revealed aligned sarcomeres and elongated mitochondria arranged between myofibrils; myofibrillar banding was evident across sections (**Fig. 2G**).

**Figure 2.**
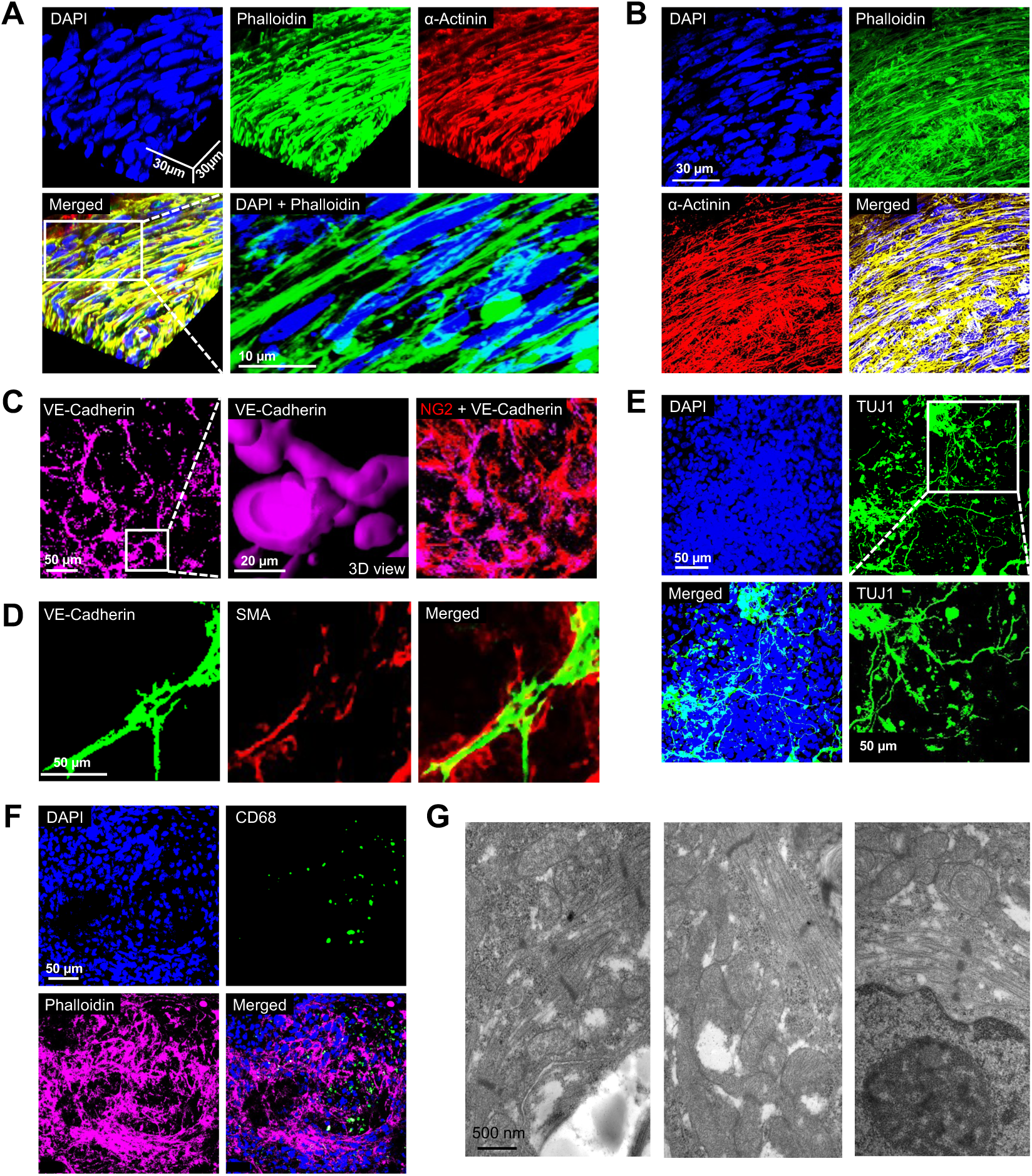
Cardiomorphs capture the cellular diversity and architecture of the human myocardium. **A-B,** CDM whole mounts stained for DAPI (blue), α-Actinin (red) and Phalloidin (green) and imaged by confocal microscopy. CDMs are presented as three-dimensional (3D) reconstruction (A) or projection (B). Representative fields are shown. Scale bar, 30 µm; inlet scale bar: 10 µm. **C,** CDM whole mounts stained for DAPI (blue), α-Actinin (green), NG2 (red), and VE-Cadherin (magenta) and imaged by confocal microscopy. Inlet depicts 3D reconstruction of vascular network and luminal structures. Representative fields are shown. Scale bar, 50 µm; inlet scale bar, 20 µm. **D,** CDM sections stained for DAPI (blue), VE-Cadherin (green), SMA (red) and imaged by confocal microscopy. Inlets depict mural cell coverage of blood vessels as projection (left inlet) and three-dimensional (3D) reconstruction (right inlet). Representative fields are shown. Scale bar, 50 µm. **E,** CDM whole mounts stained for DAPI (blue), TUJ1 (green) and imaged by confocal microscopy. Representative fields are shown. Scale bar, 50 µm. **F,** CDM whole mounts stained for DAPI (blue), CD68 (green), Phalloidin (magenta) and imaged by confocal microscopy. Representative fields are shown. Scale bar, 50 µm. **G,** Electron micrographs of mature CDMs are shown.

Cardiac maturation coincides with enhanced cardiomyocyte mitochondrial function, leading to a metabolic shift from glycolysis to glucose and fatty acid oxidation^38^. To investigate this shift, metabolomics was conducted on day 0 and day 35 CDMs. Untargeted LC-MS/MS metabolomics separated day 0 from day 35 profiles by PCA and orthogonal PLS-DA (**Extended Data Fig. 5A**)^39^. Metabolite set enrichment indicated a shift from glycolysis toward mitochondrial catabolism, including increased representation of tricarboxylic acid intermediates, acylcarnitines, and phospholipid biosynthesis pathways, with lactate and succinate levels also reduced in mature organoids (**Extended Data Fig. 4B-D**)^38,40^. Hierarchical clustering identified hyaluronan among the most increased species in later stages, consistent with its role in cardiogenesis (**Extended Data Fig. 4E**)^41^.

Cardiac metabolism is intricately linked to contractility and function^42^. To investigate if contractility patterns were altered in immature and mature CDMs, spontaneous tissue contractility was evaluated using Calcium flux analysis and IonOptix contractility measurements. These readouts corroborated cardiac maturation as evidenced by the increased transient amplitude and reduced frequency with CDM maturation, consistent with improved excitation–contraction coupling (**Extended Data Fig. 6A-F**)^42^. We next examined if CDMs replicate the physiological responses of native heart tissue to clinical compounds Phenylephrine and Carbachol. Phenylephrine enhances contractile frequency by activating ⍺-1 (ADRA1) adrenergic receptors, whereas carbachol activates Cholinergic receptor muscarinic 1 and 2 (CHRM1 and CHRM2), resulting in the redistribution and internalisation of β-adrenergic receptors B (ADRB), leading to desensitization and repressed contractile frequency^43–45^. CDMs accurately mirrored the expected contractility changes with Phenylephrine and Carbachol (**Extended Data Fig. 6G,H**)^43–45^. Additional compounds with known clinical actions, including Levosimendan, Cyclosporine, and Doxorubicin, yielded contractile and metabolic effects consistent with published pharmacology, underscoring the fidelity of the model (**Extended Data Fig. 6I-K)**^46–49^.

To explore mimicry of CDMs with native myocardial tissue, we utilised CDM integration with native mini-pig myocardium as readout. Thus, we co-cultured porcine ventricular tissue with GFP-labelled CDMs. Over 14 days, initially asynchronously beating porcine biopsies and CDMs displayed progressive attachment, absorption and integration into porcine ventricular tissue, and synchronisation of contraction by day 14 (**Extended Data Fig. 7A**). Confocal microscopy of the chimeric constructs revealed intercalation of human cardiomyocytes within porcine myocardium and anastomosis of human endothelial networks with host vasculature, as confirmed by isolectin B4 co-labelling (**Extended Data Fig. 7B,C**)^50^. Collectively, these data establish that CDMs recapitulate the multicellular composition, developmental gene programs, metabolic transitions, and physiological responses of human myocardium and can structurally and functionally integrate with native tissue.

### KSS-cardiomorphs recapitulate mitochondrial disease phenotypes

To model mitochondrial cardiomyopathy, we generated CDMs from iPSCs derived from Kearns–Sayre syndrome (KSS) patients carrying large-scale mtDNA deletions. Long-range PCR mapped deletion breakpoints to the commonly affected region spanning Complex IV and Complex I subunits, and RNA-seq confirmed reduced expression of genes within the deleted segment (**Fig. 4A-C**; **Extended Data Fig. 8A-E**). Quantitative PCR showed reduced mtDNA content relative to controls, together with decreased TFAM and PGC-1α transcripts, indicative of impaired mitochondrial biogenesis (**Fig. 4A-C**). Further, nuclear-encoded OXPHOS subunits were upregulated, consistent with compensatory mitochondrial remodelling (**Extended Data Fig. 8A-E**)^51,52^.

KSS-CDMs displayed characteristic mitochondrial mislocalisation and ultrastructural defects^53,54^. Rather than aligning between myofibrils, mutant mitochondria clustered around nuclei, producing perinuclear halos that are visible by immunostaining (**Fig. 3D**). Transmission electron microscopy also revealed swollen mitochondrial organelles with disorganised and reduced cristae density, and disrupted sarcomere patterning with shortened spacing and irregular banding, suggesting a structural and/or energetic consequence of mitochondrial impairment. (**Fig. 3E**; **Extended Data Fig. 9A**). In contrast, control CDMs exhibited elongated mitochondria with ordered cristae registering between sarcomeres, typical of healthy cardiomyocytes (**Fig. 3E**; **Extended Data Fig. 9A)** ^55,56^.

**Figure 3.**
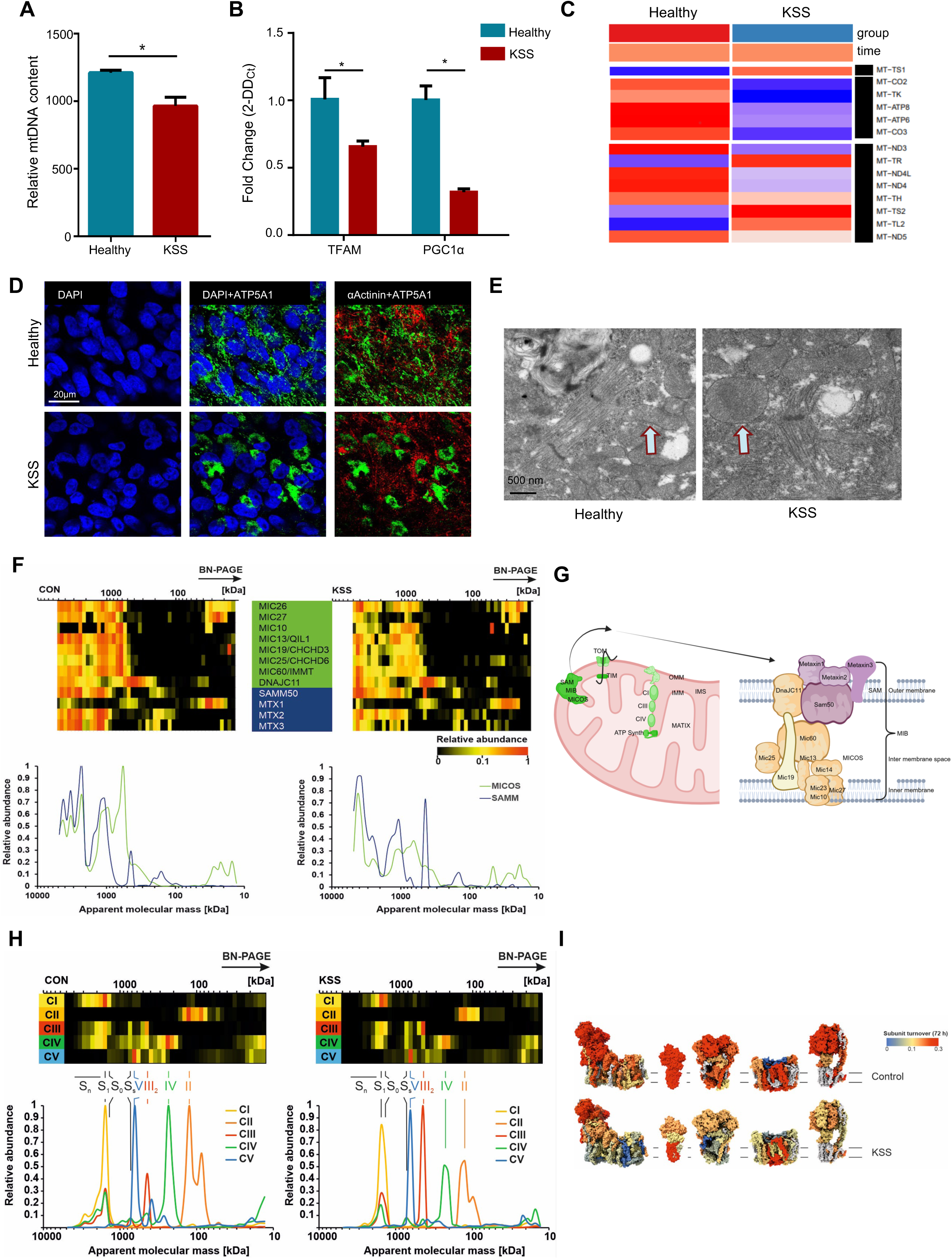
Patient-derived cardiomorphs recapitulate Kearns–Sayre syndrome (KSS) pathology. **A**, Mitochondrial DNA content in healthy and KSS CDMs (n = 3; *p < 0.05 results shown are the mean ± SEM; two-tailed unpaired t-test). **B,** Relative expression of TFAM and PGC1α in healthy and KSS CDMs (n = 3; *p < 0.05 results shown are the mean ± SEM; two-tailed unpaired t-test). **C**, RNA sequencing of mature healthy and KSS CDMs showed all mitochondrial transcripts and deleted genes. The consolidated heat maps (in z scale) were plotted from three biological replicates from each groups. **D,** Aggregated mitochondria were seen in KSS CDMs. Nuclei were stained with DAPI (blue), mitochondria were stained with ATP5A1 antibody (green) and cardiomyocytes were stained with α- sarcomeric Actinin (red). Scale bar were 20 µm. **E,** Mitochondrial ultrastructure images were taken by transmission electron microscope (TEM), showing the changes in the morphology in KSS. Scale bar was 500nm. **F,** The mitochondrial contact site and cristae organizingsystem (MICOS), which plays a crucial role in cristae architecture, is less organized in KSS organoids. Furthermore, connections with the SAM sorting and assembly machinery (SAMM) are less abundant. **G**, Schematic illustration of the MICOS and SAMM complex. **H**, The complexes of the oxidative phosphorylation system (OXPHOS) from mature KSS-CDMs exhibit fully assembled mitochondrial complexes with reductions of supercomplex organisation in complex I, III dimer and IV. **I**, Turnover of subunits within assembled OXPHOS complexes. Turnover rates upon 72 hours with medium containing heavy amino acids (pulsed SILAC) are illustrated as heatmaps within the structures of OXPHOS complexes.

**Figure 4.**
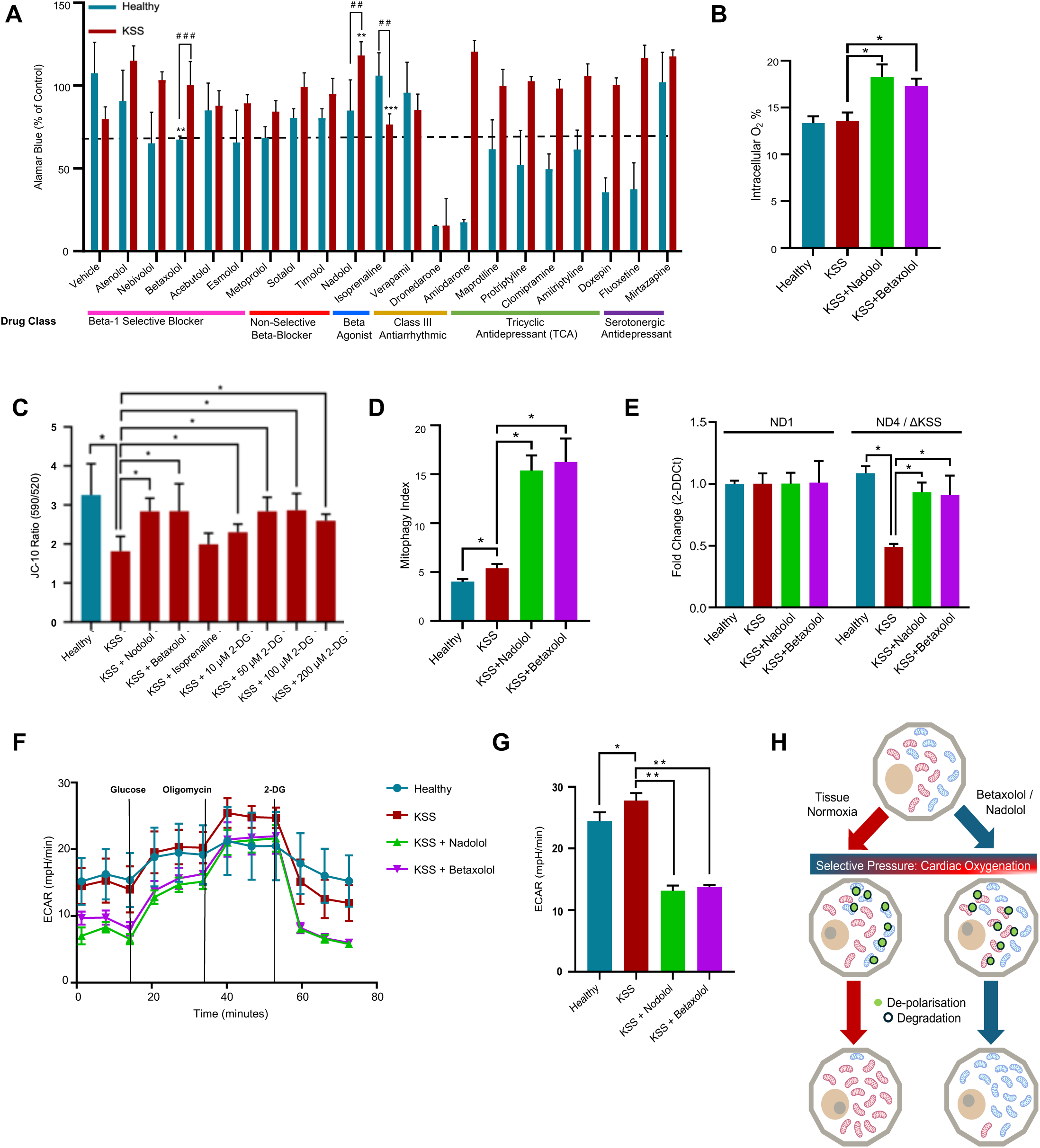
Betaxolol rebalances mitochondrial populations through dual-action quality control. **A**, Synthetic lethality screen with β-adrenergic receptor modulators to identify the drugs increasing the viability of KSS cardiomyocytes. The graph shows relative cell viability by alamarBlue assay, assessed 48 hours after treatment. The resultant data is plotted as a heatmap and expressed as a percentage (%) viability compared to vehicle control (DMSO) (n = 4 replicates per condition). **B,** Intracellular oxygenation quantification (10 wells, n=3 independent experiments) in healthy, and treated KSS CDMs. Data was analysed by multiple t test and represented as mean ± SEM. *p < 0.05. **C**, JC-10 quantification of mitochondrial membrane depolarisation (10 wells, n=3 independent experiments) in healthy, and treated KSS cardiomyocytes. Dose-escalated 2-DG treatment serves as positive control. Representative images pertaining to the analysis are shown and described in **Extended Data** Fig. 13A. Data was analysed by multiple t-test and represented as mean ± SEM. *p < 0.05. **D**, Cellular mitophagy quantification (10 wells, n=3 independent experiments) in healthy, and treated KSS cardiomyocytes. Representative images pertaining to the analysis are shown and described in **Extended Data** Fig. 13B. Data was analysed by multiple t test and represented as mean ± SEM. *p < 0.05. **E,** The ND4 gene falls within the KSS deleted region while ND1 lies at an unaffected loci, serving as an internal control. Mitochondrial DNA qPCR (with TaqMan probe) showed consistent levels of ND1 across Healthy cultures, and in untreated and Betaxolol- and Nadolol- treated KSS cultures. However, a significant reduction in ND4 levels is detected in KSS cells compared to Healthy controls (confirming the heteroplasmy in KSS cells), with ND4 levels in KSS cultures largely normalised upon Betaxolol- and Nadolol-treatment. **F**,**G,** ECAR quantification of glycolytic parameters (10 wells, n=3 independent experiments) in glucose-deprived healthy and Betaxolol- and Nadolol-treated KSS CDMs after addition of glucose (10 mM) to induce glycolysis, oligomycin (2.5 μM) to inhibit ATP synthase and 2-deoxyglucose (2-DG) (100 mM) to inhibit glucose catabolism. Data was analysed by multiple t-test and represented as mean ± SEM. *p < 0.05. **H**, Given the increase in cell viability upon Betaxolol and Nadolol treatment, we hypothesized that the suppression of oxygen consumption by the β-adrenergic antagonist Nadolol and Betaxolol may facilitate a state of increased intra-cellular oxygenation driving elevated mitochondrial function. Under these conditions, unhealthy and dysfunctional mitochondria are non-contributing and redundant, thus marked for degradation and culminating in enrichment of heathy mitochondria.

Physiologically, longitudinal respirometry revealed markedly reduced oxygen consumption in KSS-CDMs compared to controls across multi-day monitoring (**Extended Data Fig. 10A**). Seahorse-based oxygen-consumption analysis revealed significantly reduced basal respiration, blunted response to mitochondrial uncouplers and diminished spare respiratory capacity, supporting defective mitochondrial function and OXPHOS capacity (**Extended Data Fig. 10B,C**). Consistent with this, staining for JC-10, a mitochondrial membrane potential biosensor, revealed significantly reduced mitochondrial membrane potential in KSS-CDMs compared to controls, concomitant with increased glycolytic flux and reduced total ATP content, indicating a shift toward anaerobic ATP generation (**Extended Data Fig. 10C-E**). In accord, contractility assays revealed high-frequency, low-amplitude beating with increased variability in cycle timing, consistent with inefficient excitation–contraction coupling and energetic insufficiency (**Extended Data Fig. 10F**)^57^.

In assessing molecular readouts for pathological remodelling, we observed elevated expression of hypertrophic markers Natriuretic peptide (NPP)A and NPPB at the transcript concomitant to elevated levels of ACTA1, indicative of increased cardiac fibrosis (**Extended Data Fig. 11A**). KSS patients have been reported to exhibit tissue steatosis, particularly in cardiac and neural tissue^58–60^. Thus, we probed KSS-CDMs with BODIPY, a neural lipid marker for triacylglyceride (TAG) accumulation and tissue steatosis. KSS-CDMs revealed a buildup of neural lipids compared to control CDMs (**Extended Data Fig. 11B**). This observation supports the detected impairment in catabolic fatty acid oxidation and is consistent with the observed reduction in mitochondrial function in KSS-CDMs. Finally, cardiac innervation was altered in KSS-CDMs as evidenced by the reduction in TUJ1+ neurite density and neutral projection length in KSS-CDMs compared to healthy CDMs (**Extended Data Fig. 12A,B**), pointing towards impaired cardiac neurogenesis and neutral network development. Strikingly, the neurogenic defects in KSS-CDMs accurately recapitulate the neurogenic perturbations observed in KSS patient^53,54^.

To investigate possible mitochondrial structural and organisational changes as a result of the KSS mutation, we performed complexome profiling. In healthy CDMs, subunits of the mitochondrial contact site and cristae organising system (MICOS), a dynamic structure responsible for stabilising cristae junctions at the interface between the inner membrane and the inner boundary membrane, co-migrated with the sorting and assembly machinery (SAMM) in high-molecular-weight assemblies, consistent with established cristae junction scaffolds (**Fig. 3F**). However, in KSS-CDMs, high-molecular-weight MICOS–SAMM assemblies were reduced and accumulated as discrete entities with a corresponding increase in lower-mass species. These observations suggest that destabilisation of mitochondrial cristae-organising supercomplexes in KSS mutants are destabilised, and in patients with mitochondrial disorders (**Fig. 3F,G**)^61^. However, respiratory chain complexes I–V were present, indicating that gross assembly was not abolished (**Fig. 3F**). To probe mitochondrial maintenance dynamics, we used pulsed SILAC labelling to track subunit incorporation^62^. KSS-CDMs exhibited reduced incorporation across OXPHOS complexes, consistent with impaired “microregeneration” of respiratory machinery in postmitotic cells (**Fig. 3H,I**)^63,64^. These data indicate that KSS pathology in tissue involves not only mtDNA deletion-driven transcriptional loss but also supramolecular disorganisation of inner-membrane architecture and reduced capacity to replace damaged respiratory components over time (**Fig. 3I**).

Together, these findings demonstrate that KSS-CDMs recapitulate hallmark features of mitochondrial cardiomyopathy, including metabolic insufficiency, ultrastructural defects, pathological stress signalling, altered innervation, and defective maintenance of respiratory infrastructure.

### Pharmacological modulation of mitochondrial quality

We next asked whether pharmacological perturbation of oxygen handling can shift mitochondrial heteroplasmy dynamics in KSS-CDMs. The rationale derives from the observation that, in standard in vitro conditions with relatively high oxygen tension, mutant mitochondria tend to be lost over time, whereas in vivo, under lower oxygen availability, mutant heteroplasmy increases^2,65–68^. Thus, we screened US Food and Drug Administration (FDA) approved drugs known to alter oxygen consumption or cellular oxygenation, focusing on classes with clinical exposure and defined safety profiles (**Fig. 4A**). The library encompassed β-adrenergic agonists and antagonists, Class III antiarrhythmics, and antidepressant classes (tricyclic and selective serotonin reuptake inhibitors (SSRIs)). Initial viability filters in healthy CDMs removed compounds with unacceptable toxicity, which excluded most tricyclics and SSRIs at concentrations aligned with expected cellular exposure (**Fig. 4A**). Among antiarrhythmics, agents with known mitochondrial liabilities produced reductions in oxygen consumption and membrane potential in both healthy and KSS tissues, and were deprioritised. β-adrenergic agonists increased oxygen consumption but reduced KSS-CDM viability and exacerbated functional defects, consistent with an increased energetic burden under compromised OXPHOS (**Fig. 4A**).

In contrast, β-adrenergic antagonists preserved the viability of healthy CDMs and improved multiple readouts in KSS-CDMs. Within this class, the β1-selective antagonist Betaxolol and the non-selective antagonist Nadolol emerged as the most consistent positive modulators across assays (**Fig. 4A**). Both agents increased intracellular oxygenation KSS-CDMs, as measured by oxygen-sensitive probes, without inducing overt toxicity in healthy tissue (**Fig. 4B**). We then investigated the potential underlying mechanism. First, we assessed changes in mitochondrial polarisation using JC-10.

Betaxolol and Nadolol increased the fraction of polarised mitochondria in KSS cultures toward levels observed in healthy cardiomyocytes (**Fig. 4C; Extended Data Fig. 13A**). As positive control, we utilised the glycolytic inhibitor 2-deoxyglucose, which was previously shown to select against dysfunctional mitochondria^69,70^. Betaxolol and Nadolol achieved comparable improvements in shifting the fraction of polarized mitochondria towards levels present in healthy cells. (**Fig. 4C**). This led us to address mechanisms underlying the shift in mitochondrial populations. Given the lack of evidence for innate mitochondrial genomic correction, we reasoned that mitophagic events could account, in part at least, for the observed heteroplasmy shift in KSS cardiomyocytes. Thus, we co-labelled with the mitochondrial turnover sensor (MTO2) and LysoTracker to monitor mitophagosome formation and fusion with lysosomes. Both drugs increased the frequency of mitophagic events relative to untreated KSS cultures (**Fig. 4D; Extended Data Fig. 13B**). In accord, Betaxolol and Nadolol reduced the fraction of mutant mtDNA as evidenced by targeted qPCR, indicating a net shift toward wild-type organelles (**Fig. 4E**). Consequently, SeaHorse metabolic assays revealed a pronounced deficit in mitochondrial spare capacity in untreated KSS cardiomyocytes (compared to Healthy controls and KSS-treated cultures), as evidenced by the dramatic increase in ECAR levels and glycolysis upon mitochondrial ATP synthase V inhibition with oligomycin (**Fig. 4F,G**). Importantly, Betaxolol and Nadolol treatment led to an overall decrease in glycolysis and an increased reliance on OXPHOS (**Fig. 4F,G**). Together, these data indicate that Betaxolol and Nadolol drive a coupled program to increase mitophagy of depolarised organelles while concurrently driving healthy mitochondrial biogenesis to normalise and correct cellular metabolic dysfunction in KSS cardiomyocytes.

In sum, these results identify Betaxolol and Nadolol as a mutation-agnostic modulators of mitochondrial quality control that restore contractility in human cardiac tissue models of KSS. By coupling mitophagic clearance of dysfunctional mitochondria with biogenesis of functional organelles, Betaxolol and Nadolol rebalance mitochondrial populations not by targeting a specific genetic defect but by exploiting competitive fitness between organelles. Given that accumulation of defective mitochondria is a hallmark of many disorders including mitochondrial myopathies, heart disease, neurodegeneration, metabolic syndrome, and aging - this strategy may represent a generalisable therapeutic principle for restoring mitochondrial health (**Fig. 4H**).

## Discussion

The generation of cardiomorphs (CDMs) represents an advance in human cardiac organoid technology. Unlike earlier models consisting predominantly of cardiomyocytes or limited mixtures of cardiomyocytes and fibroblasts, CDMs incorporate endothelial cells, mural cells, neurons, macrophages, and stromal lineages in physiologically relevant proportions. The tissues undergo a reproducible sequence of developmental transitions, recapitulating early cardiogenesis, foetal maturation, and adult myocardial features.

Benchmarking against mouse and human developmental transcriptomes confirmed alignment at each stage. Morphological, metabolic, and functional analyses further validated the fidelity of CDMs to native myocardium. The ability of CDMs to integrate structurally and functionally with porcine heart tissue underscores their relevance as human surrogates for disease modelling and translational research. Application of CDMs to Kearns–Sayre syndrome (KSS) provided the first opportunity to study this mitochondrial disorder in a multicellular, physiologically structured human tissue context. KSS-CDMs reproduced major features of the disease, including reduced mtDNA content, loss of mitochondrial transcripts within the deleted region, compensatory nuclear upregulation of oxidative phosphorylation (OXPHOS) subunits, impaired respiration, glycolytic reliance, and diminished ATP production. Ultrastructural studies revealed mitochondrial swelling, disrupted cristae, and abnormal perinuclear clustering, consistent with findings from patient myocardium. Functional impairment in KSS-CDMs extended beyond respiration. Calcium flux analysis demonstrated contractile abnormalities characterised by high-frequency, low-amplitude cycles, reflecting inefficient excitation–contraction coupling. Metabolomic profiling confirmed shifts toward anaerobic pathways, and histological analysis revealed pathological hypertrophy and lipid accumulation. Together, these findings establish that CDMs can capture both the molecular and functional consequences of mitochondrial dysfunction. Importantly, complexome profiling provided mechanistic insight into mitochondrial structural pathology. In healthy CDMs, MICOS and SAMM complexes assemble into high-molecular-weight supercomplexes essential for maintaining cristae junctions and organising the inner mitochondrial membrane. In KSS-CDMs, these assemblies were destabilised, with accumulation of smaller subcomplexes. This observation suggests that mtDNA deletions disrupt the stability of MICOS–SAMM interactions, compromising mitochondrial architecture. While OXPHOS complexes were fully assembled, pulse SILAC experiments revealed impaired turnover of subunits, consistent with defective “microregeneration.” This phenomenon, characterised by the failure of postmitotic cells to continuously repair and replace respiratory chain components, provides a mechanistic explanation for the progressive decline in mitochondrial function in KSS.

A central observation of this study is that pharmacological modulation of oxygen consumption can influence mitochondrial selection in human cardiac tissue. In vivo, mutant mitochondria accumulate over time, whereas in standard hyperoxic culture conditions they are often eliminated. This discrepancy suggested that oxygen availability alters selective pressures on organelles. To test this, we performed a screen of approved drugs that affect oxygen metabolism. The results identified the β1-selective antagonist Betaxolol and the non-selective antagonist Nadolol as potent modulators of mitochondrial quality control. These agents reduced oxygen consumption and increased intracellular oxygenation. Mechanistic analyses demonstrated that they simultaneously induced selective mitophagy of depolarised mitochondria and promoted biogenesis of functional organelles. This dual mechanism rebalanced the mitochondrial population, reducing the fraction of mutant genomes, normalising polarization, and restoring contractility.

This approach differs conceptually from gene-targeted therapies. Rather than correcting specific mutations, Betaxolol and Nadolol exploit competitive dynamics between wild-type and mutant mitochondria. By altering intracellular oxygenation, they create an environment in which defective organelles are redundant and eliminated, while functional organelles expand. This mutation-agnostic strategy is advantageous in diseases characterized by heterogeneous or large-scale deletions for which gene correction is not feasible. The identification of β-blockers as modulators of mitochondrial quality control has immediate translational implications. Betaxolol and Nadolol are established cardiovascular drugs with well-characterized pharmacokinetics and safety profiles. Their repurposing for mitochondrial disease therefore offers a rapid path to clinical testing.

While additional studies are required to define dosing, tissue penetration, and long-term effects, the barrier to translation is lower than for novel agents.

Moreover, the mechanism of action—coupling mitophagy with biogenesis—suggests a general therapeutic principle. The accumulation of dysfunctional mitochondria is a shared feature of diverse pathologies, including cardiomyopathies, metabolic disorders, neurodegenerative diseases, and aging. Strategies that eliminate defective mitochondria while promoting functional populations may therefore have broad applicability. By demonstrating proof-of-concept in KSS, this work establishes a platform for extending investigation to other diseases. Current approaches to mitochondrial disease include gene therapy, allotopic expression of mitochondrial genes, mitochondrial replacement, and metabolic modulators. Each has limitations. Gene therapy is challenged by heteroplasmy, mitochondrial genome multiplicity, and delivery constraints. Allotopic expression and mitochondrial replacement are technically complex and have limited applicability.

Metabolic modulators such as coenzyme Q10 or dichloroacetate offer partial benefit but do not correct underlying dysfunction. The strategy identified here differs in being mutation-independent and based on pharmacological modulation of an existing pathway. Rather than targeting specific mutations or pathways, it exploits organelle-level competition. This conceptual shift is significant because it allows a single intervention to apply across multiple genetic lesions. Furthermore, because the agents are clinically approved, translation does not require de novo safety assessment. Beyond rare mitochondrial syndromes, the principle of pharmacological reprogramming of mitochondrial quality control may be relevant to common conditions. In cardiomyopathies, neurodegeneration, metabolic disease, and aging, the accumulation of dysfunctional mitochondria contributes to organ decline and impaired function. A strategy that couples selective elimination of defective mitochondria with expansion of functional populations may therefore represent a generalisable therapeutic approach.

In summary, this work establishes both a methodological advance in human cardiac organoid technology and a conceptual advance in mitochondrial therapeutics. By shifting the focus from mutation-specific interventions to organelle-level competition, it outlines a mutation-agnostic, broadly applicable strategy for diseases characterised by mitochondrial dysfunction. The translation of this principle into clinical testing may offer new treatment opportunities not only for rare syndromes such as KSS but also for more prevalent disorders of the heart, brain, metabolism, and ageing.

## Supplementary Figures (Extended Data)

**Extended Data Fig. 1 Stepwise differentiation of cardiomorphs**

**A-B**, Brightfield images of iPS cells growing in a colony form. Panel (a) shows 10x magnification, panel (b) shows 20x magnification. Scale bar, 400 µm.

**C,** iPS cells stained for DAPI (blue) and stemness markers: OCT4 (green, upper panel), NANOG (green, lower panel), SOX 2 (red). Representative fields are shown. Scale bar, 50 µm.

**D,** Brightfield image of embryonic bodies cultured in ultra-low-attachment plate. Scale bar, 200 µm.

**E,** Brightfield image of embryonic bodies cultured in a suspension. Scale bar, 400 µm.

**F,** Relative mRNA levels of stemness markers in iPSCs and CDMs at Day 0 and Day 35 of development. Levels are normalised to iPSCs mRNA . (n = 3; shown is mean ± SEM; **p < 0.01, ***p < 0.001, ****p < 0.0001; two-way ANOVA).

**Extended Data Fig. 2 Developmental timelines and benchmarking of cardiomorphs**

**A,** Schematic representation of CDM development timeline and media composition at each step of differentiation and maturation.

**B,** Representative images of CDMs at different development stages as corresponding to schematics in (a). CDMs were stained with nuclear marker (DAPI – blue), endothelial marker (ULEX – green), neuronal marker (TUJ1 – red) and cardiac marker (α-Actinin – magenta). Scale bar, 40 µm.

**Extended Data Fig. 3 Cellular composition of cardiomorphs**

**A,** UMAP plot derived from single nucleus RNA sequencing (snRNA-seq) of CDMs combined with snRNA-seq of healthy cardiac biopsies showing relative overlap of CDM-derived differentiated cell types with healthy cardiac biopsies. (n = 5022 pooled nuclei analyzed).

**B,** Quantification of cell population fractions in CDM dissociated to single cells and stained for lineage-specific antibody markers; cell types were quantified based on positive fluorescence signal and expressed as percentage (%) of total cells. (n = 3 replicates). Cell-type fractions are compared against published mean cell number values for the respective cell-types in rodent and human hearts^69–76^.

**C, D** Heatmaps showing normalized single-cell RNA sequencing (scRNA-seq) expression profiles of canonical cardiac cell-type marker genes across cell clusters derived from (**C**) human ventricular biopsy samples and (**D**) human in vitro–generated Cardiomorphs. Each column represents a transcriptionally distinct cluster, and each row corresponds to a curated panel of marker genes characteristic of major cardiac populations, including cardiomyocytes, fibroblasts, endothelial cells, smooth muscle cells, and immune cells. Expression values are Z-score–normalized and color-scaled from low (teal) to high (dark red). In human ventricular biopsies (**C**), cardiomyocyte markers (TNNT2, MYH7, ACTC1, TNNI3) and fibroblast markers (COL1A1, DCN, LUM) are prominently enriched in their respective clusters (adjusted P < 10⁻²⁰, Wilcoxon rank-sum test), validating cluster annotation accuracy. Endothelial (VWF, PECAM1) and immune-associated (PTPRC, CD68) transcripts define additional minor clusters, consistent with expected in vivo myocardial heterogeneity. In human Cardiomorphs (**D**), corresponding cell populations exhibit highly concordant expression patterns for key lineage markers (Pearson correlation r = 0.87, P < 10⁻¹⁶, compared to ventricular biopsy profiles), indicating faithful recapitulation of native myocardial architecture. Cardiomyocyte clusters display robust sarcomeric gene expression (MYL2, MYH6, TNNT2), while stromal (COL3A1, DCN) and endothelial-like (KDR, PECAM1) signatures confirm stromal and vascular mimicry within the organoid model. Limited immune-like transcript enrichment (CXCL8, CD68) suggests partial representation of inflammatory components. Collectively, comparative single-cell profiling demonstrates that human Cardiomorphs reproduce the multicellular complexity and lineage-specific transcriptomic signatures of human ventricular myocardium, with minor proportional differences in fibroblast and immune-like populations. Data represent aggregated results from n = 6 pooled donors (ventricular biopsies) and n = 3 independent Cardiomorph batches; all clusters were defined using unsupervised Leiden clustering following dimensional reduction by UMAP.

**Extended Data Fig. 4 Multi-cellularity benchmarking of cardiomorphs**

**A,** Representative images of CDMs stained with cardiac marker (α-Actinin – green) and endothelial marker (VE-Cadherin – red). Lower panel illustrates result of conversion to binary image for segmentation of CDM area. Scale bar, 200 µm.

**B,** Individual cell populations isolated from CDMs and stained with DAPI (blue) and respective cell linage markers (top panel: α-Actinin – green, SMA – red, VE-Cadherin – magenta; middle panel: Phalloidin – green, Pan Neuronal Marker – red, VE-Cadherin – magenta; bottom panel: PCM1 – green, CD68 – red, Phalloidin – magenta). Representative fields are shown. Scale bar, 100µm.

**Extended Data Fig. 5 Metabolic profiling of cardiomorph maturation**

**A,** Orthogonal projections to latent structures discriminant analysis (OPLS-DA) score plot for all metabolite features in CDMs at day 0 (D0) and day 35 (D35) (n = 4). Circles represent individual CDMs at D0 and D35 under each experimental condition.

**B,C**, Quantification of lactate (**B**) and succinate (**C**) (in ng/ml) compared between CDM at D0 and D35. Shown is mean ± SEM; *p < 0.01; two-tailed unpaired t-test with Welch‘s correction.

**D,** Metabolic set enrichment analysis (MSEA) comparing the metabolome of CDMs at D35 with CDMs at D0. The x-axis indicates the enrichment ratio calculated as the number of hits within a particular metabolic pathway divided by the expected number of hits. Highlighted pathways were significantly altered in pathway analysis with p < 0.05.

**E,** Hierarchical clustering analysis of the metabolome from CDMs harvested at D0 and D35. Depicted are metabolites with log2 (fold change) > 0.5 between CDM at D0 and D35 and with an adjusted p value < 0.01. (n = 4 replicates per group).

**Extended Data Fig. 6 Functional physiology of cardiomorphs**

**A,** Representative calcium flux transient of Day 0 and Day 35 CDM.

**B,C,** Quantification of relative amplitude (**B**) and frequency (**C**) from calcium flux transients of CDMs. (n = 3 CDMs per condition; data plotted as relative to Day 35; shown is mean ± SEM; **p < 0.01, ***p < 0.001; one-tailed unpaired t-test with Welch‘s correction).

**D-F,** Quantification of relative amplitude (**D**), departure velocity (**E**) and return velocity

(**F**) from IonOptix transients of CDMs as described in (**A**). (n = 8-10 replicates per condition; data plotted as relative to Day 35; shown is mean ± SEM; *p < 0.01, one-tailed unpaired t-test with Welch‘s correction).

**G,** Quantification of relative beating frequency from calcium flux transients of CDMs subjected to 30 minutes treatment with 200 µM Phenylephrine. (T0 = before drug addition; T30 = 30 minutes after drug addition; n = 6 CDMs per condition; data normalized to T0 and shown is mean ± SEM; ***p < 0.001; one-tailed unpaired t-test with Welch‘s correction).

**H,** Quantification of relative beating frequency from calcium flux transients of CDMs subjected to 30 minutes treatment with 0.3 µM Carbachol. (T0 = before drug addition; T30 = 30 minutes after drug addition; n = 3 iCDMs per condition; data normalized to T0 and shown is mean ± SEM; *p < 0.05; one-tailed unpaired t-test with Welch‘s correction).

**I-K,** Quantification of relative beating frequency from IonOptix transients of CDMs treated for 72 hours with 10 µM vehicle (DMSO), Levosimendan (**i**), Cyclosporine (**j**) or Doxorubicin (**k**). (n = 2-4 replicates per condition; data plotted as relative to vehicle; shown is mean ± SEM; *p < 0.01, ****p < 0.0001; one-tailed unpaired t-test with Welch‘s correction).

**Extended Data Fig. 7 Cardiomorph integration with porcine ventricular tissue**

**A,** Brightfield live imaging of integration between CDMs and adult mini pig cardiac tissue. Time snapshots at specific days are shown; black arrows point to the site of CDMs integration. Scale bar, 400 µm.

**B,** CDM-adult mini pig cardiac tissue integration block stained for nuclei (DAPI – blue), cardiomyocytes (α-Actinin – red) and human cell marker (STEM101 – green; expressed by CDMs only). Scale bar, 50 µm.

**C,** CDMs infected with pLenti-SEW-GFP (green) and integrated with mini pig cardiac tissue. Organoid-mini pig cardiac tissue fusions were subsequently stained for nuclei (DAPI – blue), endothelial cells (IB4 – red), and cardiomyocytes (α-Actinin – magenta). White line in inlet points to endothelial integration of CDM with rat tissue. Scale bar, 50µm; inlet scale bar 25µm.

**Extended Data Fig. 8 Compensatory gene expression in KSS**

**A-E,** RNA sequencing of healthy and KSS CDMs mitochondrial ETC complex transcripts and deleted genes (in black squares). The consolidated heat maps (in z scale) were plotted from three biological replicates from each groups.

**Extended Data Fig. 9 Perturbed mitochondrial organization and patterning in KSS**

**A,** Healthy and KSS iPSCs were differentiated into cardiomyocytes (CM). In CMs, nuclei were stained with DAPI (blue), mitochondria were stained with ATP5A1 antibody (majenta) and cardiomyocytes were stained with α- actinin (red). Scale bar were 20 µm.

**Extended Data Fig. 10 Energetic and functional deficits in KSS cardiomorphs**

**A**, Healthy and KSS CDM OCR was measured by Resipher system from Lucid scientific. After the real time monitoring of the OCR for 72 hrs (10 wells, n=3 independent experiments), the data was represented as mean ± SEM.

**B,** Mitochondrial OCR of healthy and KSS CDMs were measured by Seahorse XF96 analyser by using Palmitate as a substrate. The result was normalized with total cell number per well (10 wells, n=3 independent experiments). OCR was measured under basal conditions and after addition of oligomycin (2.5 µM) to inhibit ATP synthase, FCCP (5 µM) to uncouple oxidative phosphorylation and antimycin A (2.5 µM) and rotenone (2.5 µM) to measure non-mitochondrial respiration. Data was represented as mean ± SEM. Data was analysed by multiple t test and represented as mean ± SEM. *p < 0.05.

**C,** Healthy and KSS CDMs were stained with JC-10 dye and the graph was plotted after taking the reading in plate reader with JC-10 green and JC-10 red channel. Data was analysed by multiple t test and represented as mean ± SEM. *p < 0.05.

**D**, ECAR quantification of glycolytic parameters (10 wells, n=3 independent experiments) in glucose-deprived healthy and KSS CDMs after addition of glucose (10 mM) to induce glycolysis, oligomycin (2.5 μM) to inhibit ATP synthase and 2-deoxyglucose (2-DG) (100 mM) to inhibit glucose catabolism. Data was analysed by multiple t-tests and represented as mean ± SEM. *p < 0.05.

**E,** Total ATP content of healthy and KSS CDMs was measured (n=3 independent biological replicates) by the EnzyLight™ ATP Assay Kit by the manufacturer’s protocol. Data was analysed by multiple t test and represented as mean ± SEM. *p < 0.05.

**F**, Representative calcium flux transients assay of healthy and KSS CDMs. (n=3 independent biological replicates).

**Extended Data Fig. 11 KSS cardiomorphs exhibit elevated pathologic hypertrophy marker expression**

**A,** Relative mRNA levels of cellular stress markers in healthy and KSS CDMs. n = 3 replicates (biologically independent samples). Statistical analysis was done by multiple t-test. All bar graphs show mean ± SEM. *p < 0.05.

**B,** Healthy and KSS iPSCs were differentiated into cardiomyocytes (CM). In CMs, nuclei were stained with DAPI (blue), mitochondria were stained with NPPA antibody (majenta), cardiomyocytes were stained with α- actinin (red) and BODIPY (green), for neutral lipid detection. Scale bar were 20 µm.

**Extended Data Fig. 12 Dysfunction cardiac neurogenesis in KSS cardiomorphs**

**A,B,** KSS CDMs showed less neuronal density and projection compared to the healthy one. Nuclei were stained with DAPI (blue), neurons were stained with TUJ1 antibody (green) and cardiomyocytes were stained with α-Actinin antibody (red). Scale bar was 50 µm.

**Extended Data Fig. 13 KSS mitochondria exhibit membrane depolarization and autophagic tagging**

**A,** Representative images of Healthy and KSS cardiomyocytes treated with Betaxolol or Nadalol, respectively, stained with the JC-10 membrane depolarisation dye and for ATP5A1, as marker for both healthy and mutant mitochondria, and analysed by confocal microscopy.

**B,** Representative images of Healthy and KSS cardiomyocytes treated with Betaxolol or Nadalol, respectively, stained with Lysosomal and Mitophagy markers, as readout for Mitophagy, and analysed by confocal microscopy.

## Acknowledgements

We thank P. Schuster, N. Ivanovski, J. Maxeiner, H. Fischer, J. Meisterknecht, all members of the Institute for Cardiovascular Regeneration and Genome Biologics for technical, regulatory, administrative, analytical, reagent and scientific support. This work was supported by the LOEWE Center for Cell and Gene Therapy and the European Innovation Council (GA: 822455) to J.K., the SFB-TRR 267 (Non-coding RNA in the cardiovascular system, #403584255), SFB1531 (Damage control by the stroma-vascular compartment, grant #456687919) to S.D. and I.W, and the German Research Foundation (DFG) (Exc2026) to S.D. and J.K., and instrument grant support (INST 515/28-1 FUGG) to M.P., the European Research Council (Angiolnc) to S.D., and the Foundation for Pathobiochemistry and Molecular Diagnostics to P.M. Y.W. was supported by the China Scholarship Council (CSC) Grant #202108080020. I.W. was supported by DFG (grant #515944830).

## Author Contributions

M.D.P., Y.W., D.S. and J.K. planned and designed the experiments, and wrote the paper with M.D., I.W. and J.W. M.D.P. and Y.W. executed most of the experiments with experimental and analytical help from T.Y., A.P.D., M.W., C.Z., T.A., P.G., T.T.D., J.Kaur and N.I.. I.W. and A.C-O. performed complexome profiling analysis and illustrations. M.L., M.P. and P.M. performed metabolomics studies and data analysis; L.N. and W.A. performed scRNAseq studies and analyzed the data. A.Z. and S.D. provided supporting data, scientific and technical expertise.

## Supporting Online

### Material Materials and Methods

#### iPS cells maintenance

Human induced pluripotent stem cells (hiPSCs) line WSTIi-81A (Merck, 66540196) was cultured at 37°C / 5% CO_2_ in complete TeSR^TM^-E8^TM^ media (STEMCELL^TM^ Technologies, 05990), according to supplier’s instruction and with addition of 1% Penicillin/Streptomycin (Sigma-Aldrich, P4333-100ml). Prior to plating, iPSCs culture dishes were pre-coated with iMatrix 511 (Takara, T304) for 1 hour at 37°C. For maintenance, media was changed daily. KSS iPSC^71^ (with 6kb mtDNA deletion) were kindly provided by Oliver Russell (Newcastle university, UK). KSS iPSC were maintained on Matrigel coated dishes with NutriStem hESC XF medium (Biological Industries cat, 05-100-1 A) supplemented with 1× sodium pyruvate and 0.05 mg/ml uridine. For passaging, cells were dissociated by ReLeSR reagent and replated at a density of 1.5 × 104 cells/cm2 in NutriStem with 10 µM Rock inhibitor (Y-27632) (Stemcell Technologies, 72302). Fresh NutriStem media was replenished after 24 hrs. Cells detachment was performed with ReLeSR (STEMCELL^TM^ Technologies, 05872) for 10 to 15 minutes incubation at 37°C / 5% CO_2_. Cells were harvested gently with complete TeSR^TM^-E8^TM^ media, supplemented with 10 µM ROCK-inhibitor (STEMCELL^TM^ Technologies). Medium with ROCK-inhibitor was replaced with complete TeSR^TM^-E8^TM^ media after 24 hours of incubation.

### Mini pig cardiac explants

Adult Göttingen mini-pig cardiac live biopsies were kindly provided by Lars Friis Mikkelsen (Ellegaard Göttingen Minipigs A/S).

### CDM formation and maintenance

hiPSCs were plated (500 cells per well) in AggreWell^TM^ 800 plate (STEMCELL^TM^ Technologies, 34821) at Day −10 (per timelines schematics presented in **fig. S2a**). Cells were cultured for 2 days in TeSR^TM^-E8^TM^ media at 37°C / 5% CO_2,_ to form embryonic bodies (EBs). TeSR^TM^-E8^TM^ medium was sequentially switched to differentiation Medium 1 (medium composition in **table S2**; Day −8) and Medium 2 (medium composition in **table S2**; Day −6). At Day −4, differentiation medium was replaced with Medium 3 (medium composition in **table S2**) and refreshed once every 2 days. At Day 0, Medium 3 was replaced with maintenance Medium 4 (medium composition in **table S2**), at which point EBs were fully differentiated to CDMs. Medium 4 was refreshed every second day for 10 days, subsequently being replaced with Medium 5 (medium composition in **table S2**). CDMs were kept in Medium 5 for 4 days with media change every second day, followed by final replacement to Medium 6 (medium composition in **table S2**). CDMs were maintained in Medium 6 with media change every second day until ready for further assays.

### CDM dissociation assay

Per sample, 40 CDMs were collected in 1.5 mL tube, and 1 mL 2.5% Trypsin (Merck, 59427C- 100ML) was added. The tube was centrifuged at 1400 rpm for 15 minutes at 37°C (Eppendorf Thermomixer Compact Next). The cell pellet was pipetted up and down several times until most cells were dissociated and then centrifuged for another 5 minutes. 1 mL of culture medium (Medium 6; medium composition in **table S2**) was added to the tube to further stop the dissociation.

### Synthetic lethality screen

iPSC-derived wildtype and KSS mutant cardiomyocytes were seeded in 384-well plates with 2500 cells / well (Greiner Bio-One, 781186). Cells were subsequently treated with compounds from the FDA Approved Compound Library (Targetmol), that were added at 10 μM; 1% (10 µM) DMSO (Carl Roth, A994.2) was used as a solvent (vehicle) control. 72 hours later cell viability was measured using Alamar Blue assay (Bio-rad, BUF012B).

### Trypan blue assay

Cell viability was measured by mixing 0.4% Trypan blue solution (Gibco, 15250061) with equal volume of myeloid cells suspension, following by counting the live (transparent) and dead (blue) cells under the microscope.

#### Calcium transient assay

CDMs were collected into 2 mL eppendorf tube and incubated with labelling medium (Cal-520® 2.5 µM and Pluronic® F-127 0.04% in PBS) for 90 minutes at 37°C / 5% CO_2_. After the labelling process was finished, the CDMs were washed twice with PBS and once with pre-warmed Plain Medium 6 (Medium 4 mixed with Medium 2; Medium 2 and Medium 4 composition in **table S2)**. CDMs were then distributed into 384-well plate (one CDM per well) with ultra-low-attachment surface and black walls (Corning, 4516). 100 µL of Plain Medium 6 was added and CDMs were further incubated for 90 minutes at 37°C, 5% CO_2_ in a Plate Reader (ClarioStar, 430-0911) before the measurement. The fluorescence intensity was measured 750 times with the interval of 0.02 s between the reading points.

For drug treatments, drugs were prepared at the concentration of 100x with 1 µL per well added for the drug treatment to reach final concentration of 10 µM. CDMs fluorescence intensity was measured before drugs addition and then following minimum of 30 minutes drug treatment.

### Resipher measurement

Single CRM was put into single well of low attachment U bottom 96 well plate with the CRM maintenance medium. Afterwards the sensor was connected to the lid and the sensors were dipped into the media. This setup was kept inside the cell culture incubator and it was connected to its cloud software (Lucid lab) via WLAN connection. The OCR measurement was recorded for 72 hours and analyzed by the same software.

### Seahorse measurement

Basal Assay medium and rest XF96 consumables were purchased from Seahorse Bioscience Inc. CDMs were kept in Poly-D-Lysine pre coated Seahorse plate for 4 hours at 1 CDM/well. OCR was measured using Palmitate as a substrate in the XF96 basal assay medium. OCR was measured under basal conditions and after addition of oligomycin (2.5 µM) to inhibit ATP synthase, FCCP (5 µM) to uncouple oxidative phosphorylation and antimycin A (2.5 µM) and rotenone (2.5 µM) to measure non-mitochondrial respiration. ECAR was measured as per the Seahorse XF Glycolysis Stress Test Kit protocol by subsequent injection of Glucose (10 mM), Oligomycin (2.5 mM) and 2-DG (50 mM). After the measurement, the data was normalized by the total cell number measured by Alamar Blue assay (Bio-rad, BUF012B).

### JC10 measurement

JC-10 (ENZ-52305) was used to determining mitochondrial membrane potential of healthy and KSS CRMs in microplate-based fluorescent assays. JC-10 was added to the CRM maintenance medium (15 µM working concentration) for 1 hour at 37℃ incubator. After one hour of incubation, the JC10 was washed by PBS and the reading was taken in the microplate reader in green and red channel.

### ATP concentration measurement

Healthy and KSS CRM ATP content was measured by the EnzyLightTM ATP Assay Kit (EATP- 100) by the manufacture’s protocol. The ATP concentration was determined from a standard graph.

### cTnT assay

Cells or CDMs pre-conditioned cell culture media were collected and then diluted to 1:3 with NS Reagent from the cTNT Elisa kit (Abcam, ab223860). cTNT was evaluated by the cTNT Elisa kit following the manufacturer’s instructions. Fluorescence was read out with ClarioStar plate reader (430-0911).

### Fluorescence immunohistochemistry

CDMs were collected and fixed in 4% PFA (Santa Cruz, CAS 30525-89-4) for overnight at 4oC. For whole-mount staining, fixed CRMs were permeabilized with 1% Triton X-100 in PBS for 20 mins, followed by blocking with 5% horse serum (Sigma-Aldrich, H0146) in PBS for 1 hour at room temperature. Primary antibodies (**table S3**, 1:200 anti-α sarcomeric actinin, #A7811, Sigma Aldrich, 1:200 anti-NPPA, #27426-1-AP, Proeintech, 1:200 anti ATP5A1, #ab14748, 1:200 anti-alpha smooth muscle Actin, # ab202295, 1:200 anti- Anti-beta III tubulin(Tuj1), #ab18207) were diluted in 2% horse serum with 0.002% Triton X-100 and incubated overnight at 4°C. Afterwards samples were washed for 3 times with PBS containing 0.002% Triton X-100 for 15 minutes each at room temperature. Secondary antibodies (1:500 AlexaFlour 488, #A11017, 1:500 AlexaFlour 555, #A21425 and 1:500 AlexaFlour 647,# A-21245) were diluted in 2% horse serum with 0.002% Triton X-100 and incubated for 3 hours at room temperature in the dark. After three additional washes (for 15 minutes each at room temperature), CRMs were mounted using ProLong Gold Antifade Mountant (Thermo Fisher Scientific, P36934) and imaged with a Leica TCS SP8 confocal microscope.

### Fluorescence quantification

Apoptosis and infiltration quantification was performed by analyzing the area covered by positive Cleaved-Caspase-3 and Qtracker signal respectively (ImageJ). The measurements were acquired from maximum Z-projections of 3 independent and non-overlapping z-stacks per SCO. Equal intensity threshold was set for all images as a cut-off for binary conversion. The marker positive area was segmented on binary images and expressed as a fraction of the total area (set equal for conditions). For average area quantification a mean value of all areas per condition was quantified. Fibrosis quantification was performed by analyzing mean intensity of COL1A1 from Z-projections of 3 independent and non-overlapping z-stacks per each SCO after background subtraction.

### Electron microscopy

CRMs were perfused with 1.5% glutaraldehyde and 1.5% paraformaldehyde in 0.15 M HEPES buffer. Afterwards, CRMs were removed and stored in the same fixative at 4°C for at least 24 h. For epoxy resin embedding, the post fixed CRMs were fixed in 1% osmium tetroxide in aqua bidest, stained en bloc in half-saturated watery uranyl acetate, dehydrated in an ascending ethanol series and finally embedded in Agar 100. Ultrathin sections were cut using an ultramicrotome and examined by TEM (Zeiss EM902).

### Mitochondrial complexome profiling

Raw data of mass spectrometry, a detailed description of material and methods and extended data analysis including complexomes and turnover rates have been deposited to the ProteomeXchange Consortium via the PRIDE partner repository with the dataset identifier PXD063313 and PXD 063322111. Reviewer account details: Complexome (Login here: https://www.ebi.ac.uk/pride/profile/reviewer_pxd063313, Reviewer account details, Username, Password: uCW8l4qs9upJ, Go to “ Review submission” and Turnover rates (Login here: https://www.ebi.ac.uk/pride/profile/reviewer_pxd 063322, Reviewer account details, Username: reviewer_pxd, Password: 1yc5jUfBDinX, Go to “ Review submission”.

### Lentiviral GFP expression

Lentiviral particles were generated by transfecting HEK293T cells with the packing plasmids pCMVΔR8.91 and pMD2.G as well as the GFP-ORF carrying plasmid SEW. In brief, HEK293T cells were cultured in a T175 flask (Greiner Bio-One) until reaching a confluency of 70%. A transfection mix of 100 µL Opti-MEM (Life Technologies) containing 6 µg of pCMVΔR8.91, 2 µg of pMD2.G and 8 µg of SEW was prepared that pre-incubated for 20 minutes at room temperature before adding to the HEK293T culture. After 4 days of culturing, culture supernatant was cleared from cellular debris by centrifugation (500x g, 5 minutes, 4°C) and concentrated 40x using the Amicon Ultra-15 centrifugal units 100 KD (4 times, 2500x g, 30 minutes, 4°C).

For GFP expression, CDMs were distributed one per well in a 96-well plate with ultra-low- attachment surface (Corning) and virus-containing supernatant was titrated on CDMs for 24 hours to determine the titers needed to transduce >95% of the cells. The virus supernatant was then replaced with Medium 6 (medium composition in **table S2**) and 50% of medium was changed every second day for 3 days. After that, GFP expression was confirmed by fluorescence microscopy.

#### Metabolomics Analysis by LC-MS/MS metabolite profiling

Metabolite analyses based on liquid chromatography tandem mass spectrometry (LC-MS/MS) were performed using a targeted approach for profiling of tricarboxylic acid cycle (TCA) metabolites including lactate as well as untargeted metabolomics profiling. TCA cycle metabolite profiling was done as described elsewhere ^72^. In brief, sample extracts were supplemented with internal standard used for quantification of TCA cycle metabolites, followed by a dry-down using vacuum-assisted centrifugation. Residues were reconstituted in initial mobile phase conditions and immediately analyzed by LC-MS/MS. Leftover sample materials were dried down, residues were reconstituted in 200µL aqueous acetonitrile (10% water) including 0.1% formic acid and transferred into auto sampler glass vials. A pool consisting of 20 µL from each of those samples was prepared in a separate autosampler vial and was analyzed multiple times but randomly throughout the analysis. Samples were analyzed by liquid chromatography high resolution tandem mass spectrometry (LC-HR-MS/MS) using an instrument set-up containing an ultra-performance liquid chromatography system (Aquity I-class, Waters) coupled to a quadrupole – time-of-flight (QToF) additionally equipped with an ion mobility separation device (IMS) (VION IMS QToF, Waters). Chromatographic separation was achieved by using an XBridge BEH Amide column (2.1 x 100mm, 1.7µm, Waters) at 45°C using a gradient of mobile phase A (water including 0.1% formic acid) and mobile phase B (acetonitrile including 0.1% formic acid). After sample injection starting with 1%/99% A/B at a flow rate of 0.4 mL/min, mobile phase A was linearly increased to 5% within 2 minutes, followed by a further increase to 99% reached at 7.5 minutes. After 0.5 minutes, starting conditions were re-established within 0.3 minutes followed by column equilibration at starting conditions for 1.7 minutes. Positive and negative electrospray ionization (ESI) in combination with a high definition data acquisition scan mode (HDMS^E^) that included ion mobility screening in conjunction with determination of metabolite specific collision cross section (CCS) values, and accurate mass as well as respective fragment ion mass screenings. The observed mass range included mass-to-charge ratios between 50 and 1000 Da. Total scan time was set to 0.3 s from which 40% of the time a collision energy of 6eV (low energy) was applied before collision energy was respectively ramped either from 15 to 50 eV or 10 to 80 eV (high energy) in positive or negative ESI mode for the remaining 60% of the time. Ion source parameters in positive ESI mode were respectively set to 1kV and 40V for capillary voltage and sample cone voltage and to 120°C and 550°C for source and desolvation temperature. In negative ESI mode, parameters were set to 1.5kV and 40V, and to 120°C and 450°C, respectively. Nitrogen was applied as cone gas and desolvation gas with flow rates of 50 L/h and 1000 L/h, respectively, in positive and negative ESI. Repeated injections of a Leucine-Enkephalin solution served as reference for mass correction. Data acquisition and raw data processing was performed by using the Waters Unifi software package 2.1.2. For further data processing, Unifi export files derived from positive and negative ESI screenings were imported into the Progenesis QI software (Nonlinear Dynamics, Newcastle upon Tyne, UK). Processing included peak picking with auto threshold and chromatographic alignment using an automatically chosen reference sample, signal deconvolution and data normalization as well as set-up of experimental design. For metabolite identification, features derived from untargeted metabolomics screenings were compared with the publicly available human metabolome database (HMDB, www.hmdb.ca) using the Progenesis Metascope plugin applying a precursor mass accuracy of ≤ 10 ppm and a theoretical fragment mass accuracy of ≤ 10 ppm. Furthermore, features were compared to in-house data achieved by injections of metabolite standards in neat solution including allowed deviations of retention time and CCS of 0.3 minutes and 3%, respectively. Analysis derived features that have not received a metabolite identification were excluded, remaining features in conjunction with their identifications were exported to Microsoft Excel for further data processing.

### Tissue integration

The adult rat heart was isolated and soaked in ice-cold sterile 0.02M BDM (2,3-Butanedione monoxime) in HBSS buffer (Gibco). The ventricular tissue was then sectioned into small blocks and maintained in medium 6 in ultra-low-attachment 96-well-plate overnight at 37°C, 5% CO_2_. The next day, GFP-labelled CDMs were transferred into the well with tissue for co-culture. Medium 6 (Medium 6 composition in **table S2**) was changed daily. The integrated CDM-tissue were then collected, fixed with PFA 4% for 24 hours and cryo-sectioned (50 µm thickness). The sections of tissue-CDM were used later for immunofluorescence staining.

### RNA isolation, reverse transcription and qRT-PCR

Samples were harvested and the total RNA was isolated using the RNeasy Mini Kit (Qiagen, 74106), as per manufacturer protocol. RNA was reverse transcribed into cDNA using RNA to cDNA EcoDry Premix (random hexamers) kit (Clontech, Cat No 639545) following the manufacturer’s instructions. Target genes were quantified with RT-qPCR using Fast SYBR™ Green Master Mix (#4385612, Thermo Fischer scientific) in an ViiA™ 7 Real-Time PCR System with 384-Well Block PCR machine (Thermo Fischer scientific). Target genes were normalized by HPRT. Relative expressions of the genes were calculated by ΔΔCt method. Primers used are listed in **table S4**.

### RNA-seq data acquisition

Mouse RNA sequence data were downloaded from BioProject PRJNA306360 (E8.5), BioProject PRJNA66167 (E14.5), BioProject PRJNA489472 (P1), BioProject PRJNA378351 (P56). The genes of mouse and CDMs are filtered from Heart Development (GO0007507), and the gene counts are normalized by log2 (counts/million+4). The correlation coefficient is calculated by the Pearson correlation coefficient. The transcriptomic datasets of mouse heart tissue at different developmental stages were first downloaded from BioProject PRJNA306360 (E8.5), BioProject PRJNA66167 (E14.5), BioProject PRJNA489472 (P1), BioProject PRJNA378351 (P56). Raw sequencing data were processed with Trimmomatic, aligned to the Ensembl mouse genome (GRCm39) with STAR, and mapped with HTSeq. By default, a filter is also applied to remove genes expressed at the level of 1 in samples for normalized expression data. After removing outlier samples by exploratory hierarchical clustering, at least three replicates are available at each stage of development. The orthologous gene we used to calculate cardiomyocyte developmental tendency between corresponding stages of the human-induced Pluripotent Stem Cells and mouse are from the Heart Development (GO0007507). Human and mouse quantile-normalized gene-expression data were integrated using the Seurat pipeline, the counts in the matrix were normalized and natural log transformed by the default methods of Seurat. The data were then scaled using the standard Z-score scaling before Pearson correlation analysis.

### Human single-nucleus, single-cell RNA-sequencing data and processing

Two human sn/scRNA-seq datasets were used: data from healthy cardiac tissue from the left ventricle of 21 individuals in the Litvinukova et al. study^20^ and data from CDMs derived in Frankfurt. For healthy cardiac biopsy tissue processing, see original publication^20^. For CDM nucleus isolation, the following protocol was used: CDMs were pre-digested with a 5-ml enzyme solution of collagenase (2,500 U, Thermo Fisher Scientific) in HBSS+/+ (Gibco) for 10 min at 37 °C in a water bath. After centrifugation at 500*g* and 4 °C for 5 min, the supernatant was discarded, and nuclei were isolated after cell disruption with a glass dounce homogenizer (five strokes with a loose pestle and ten strokes with a tight pestle). After filtering (20-µm strainer, pluriSelect), the suspension was centrifuged at 1,000*g* and 4 °C for 6 min and resuspended in 500 µl staining buffer containing 1% BSA (Sigma-Aldrich), 5 nM MgCl_2_ (Sigma-Aldrich), 1 mM EDTA (Gibco), 1 mM EGTA (Gibco), 0.2 U µl^−1^ RNasin Plus Inhibitor (Promega) and 0.1 µg ml^−1^ Hoechst (Life Technologies) in Dulbeccós phosphaste buffered saline (DPBS). Hoechst-positive nuclei were separated from cell debris by using the FACSAria Fusion instrument (BD Biosciences) and sorted into staining buffer without Hoechst at 4 °C.

### Single-nucleus RNA-sequencing library preparation

CDM nuclear suspensions were loaded on a 10x Chromium Controller (10x Genomics) according to the manufacturer’s protocol. All snRNA-seq libraries were prepared using the Chromium Single Cell 3′ version 3 Reagent kit (10x Genomics) according to the manufacturer’s protocol. Individual nuclei were isolated into droplets together with gel beads coated with unique primers bearing 10x cell barcodes, UMIs and poly(dT) sequences. Reverse transcription reactions generated barcoded full-length cDNA followed by disruption of emulsions using the recovery agent and cDNA cleanup with DynaBeads MyOne Silane Beads (Thermo Fisher Scientific). Total cDNA was amplified using a Biometra Thermocycler TProfessional Basic Gradient with a 96-well sample block (98 °C for 3 min; cycled 14×: 98 °C for 15 s, 67 °C for 20 s and 72 °C for 1 min; 72 °C for 1 min; held at 4 °C). Amplified cDNA products were cleaned with the SPRIselect Reagent kit (Beckman Coulter). Indexed sequencing libraries were constructed using reagents from the Chromium Single Cell 3′ version 3 Reagent kit as follows: fragmentation, end repair and A tailing; size selection with SPRIselect; adaptor ligation; post-ligation cleanup with SPRIselect; sample index PCR and cleanup with SPRIselect beads. Library quantification and quality assessment were performed using the Bioanalyzer Agilent 2100 with a High Sensitivity DNA chip (Agilent Genomics). Indexed libraries were pooled equimolarly and sequenced on the Illumina NovaSeq 6000 by GenomeScan using paired-end 26 × 98-bp reads as the sequencing mode.

### Single-nucleus RNA-sequencing data analysis

CDM single-nucleus expression data were processed using the Cell Ranger Single Cell Software Suite (version 3) to perform quality control, sample de-multiplexing, barcode processing, and single-cell 3′ gene counting. Sequencing reads were aligned to the human reference genome GRCh38. Further analysis was performed in Seurat (v3.1) R (v4). Data were filtered based on the number of genes detected per cell. A global-scaling normalization method for the gene-cell-barcode matrix of the samples was employed. We normalized the feature expression measurements for each cell by the total expression, multiplied this by a scale factor (10,000), and log-transformed the result to yield the normalized unique molecular identifier (nUMI) value later reported. Regression in gene expression was performed based on the number of normalized unique molecular identifiers (nUMI). PCA was run on the normalized gene-barcode matrix. The CDM dataset was analysed alone and after merging with the cardiac biopsy dataset in Seurat. Differential expression was performed using the FindMarkers and FindAllMarkers tools using a Bonferroni adjusted p-value less than 0.05 as a significance threshold.

### Statistical analysis

Statistical analysis was performed using GraphPad Prism 9 software. Statistical analysis for pairwise comparisons was performed via two-tailed t-tests with Welch’s correction. Data with more than 1 variable was evaluated by two-way ANOVA followed by multiple comparisons tests (with Dunnett’s or Sidak’s corrections, as appropriate). A *p*-value of less than 0.05 was considered statistically significant. No statistical methods were used to predetermine sample size.

