## Extended Data Figures 1-13 for "Human heart organoids reveal a regenerative strategy for mitochondrial disease"

#### Extended Data Fig. 1

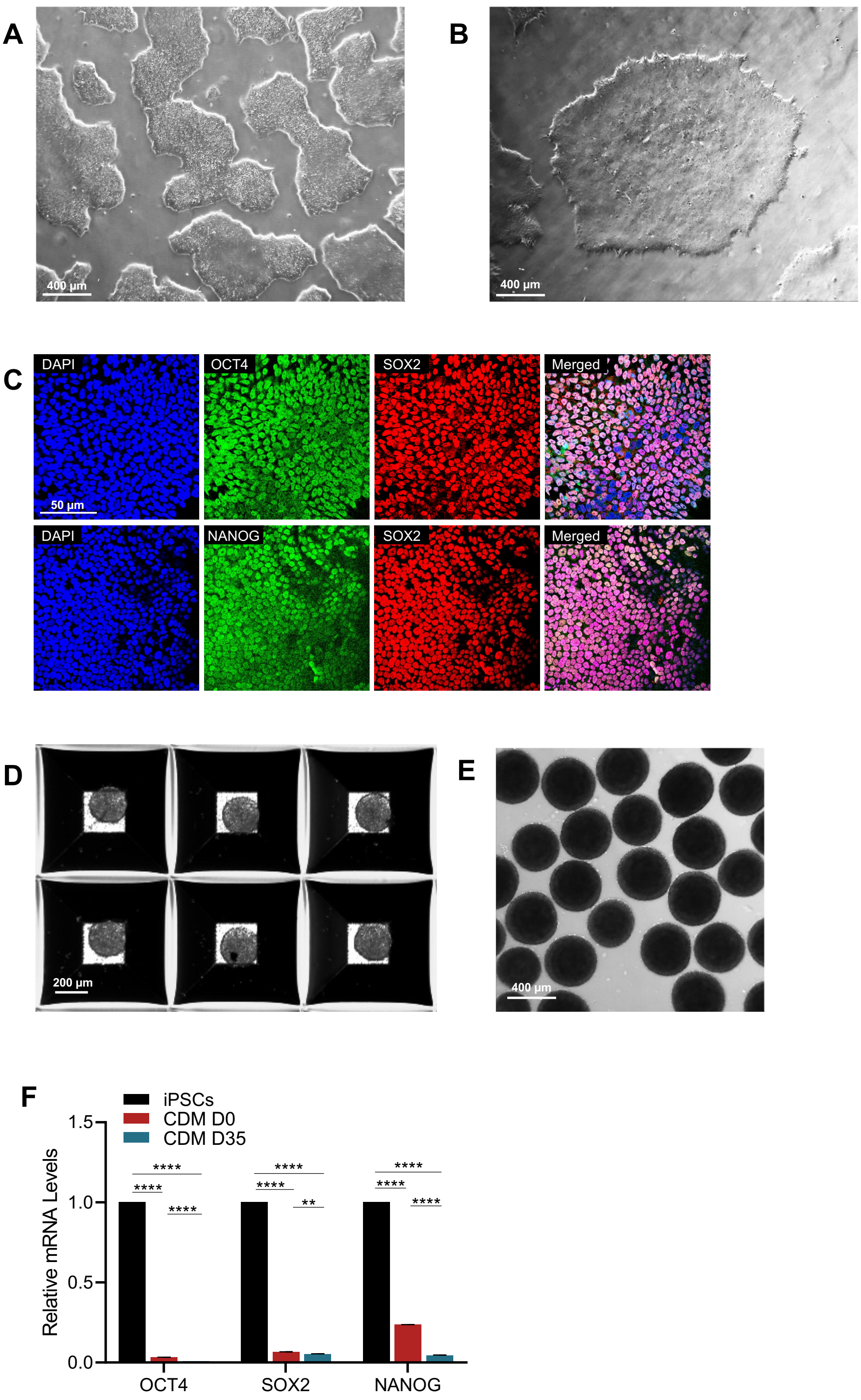

#### Extended Data Fig. 2

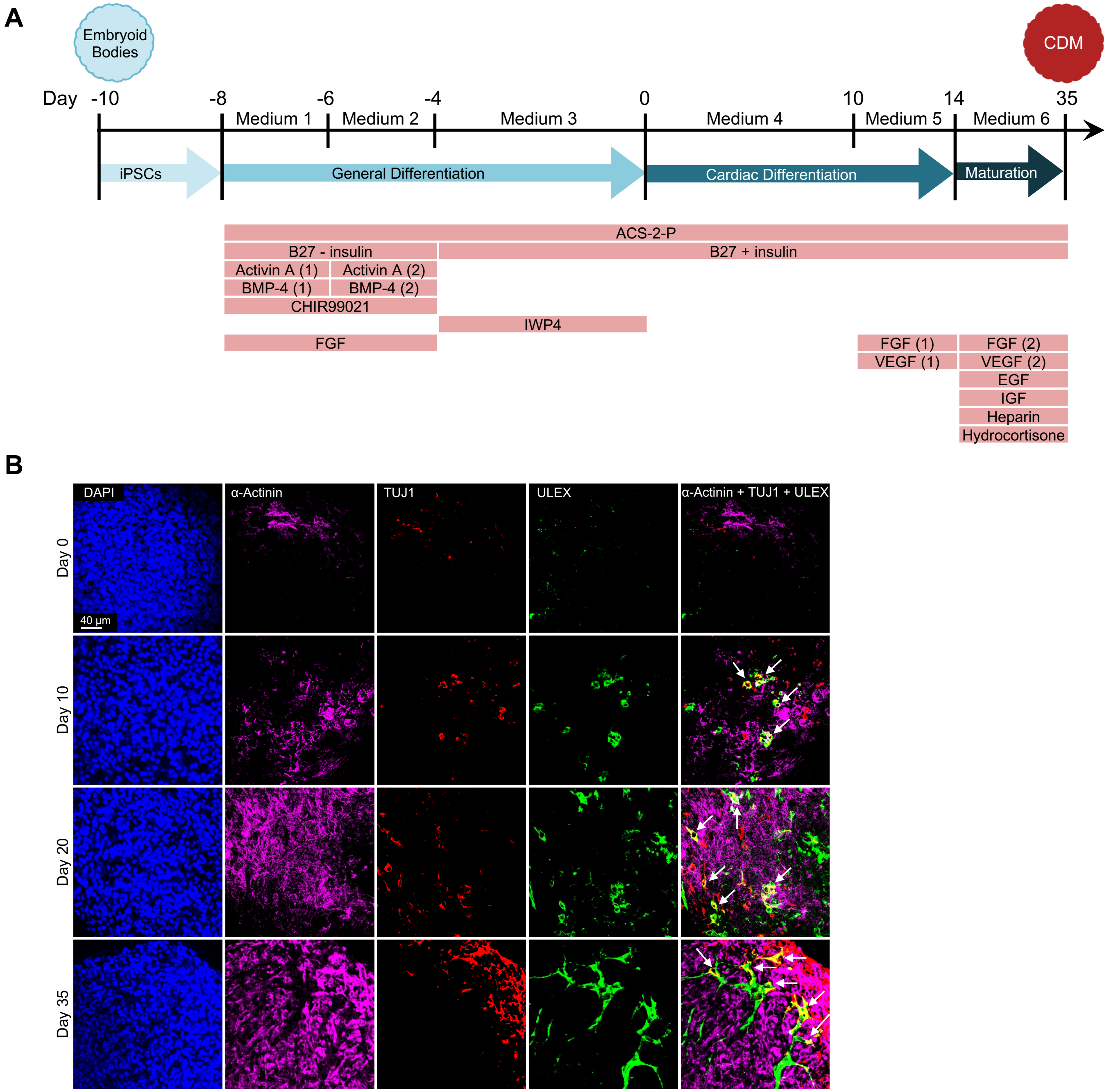

### Extended Data Fig.3

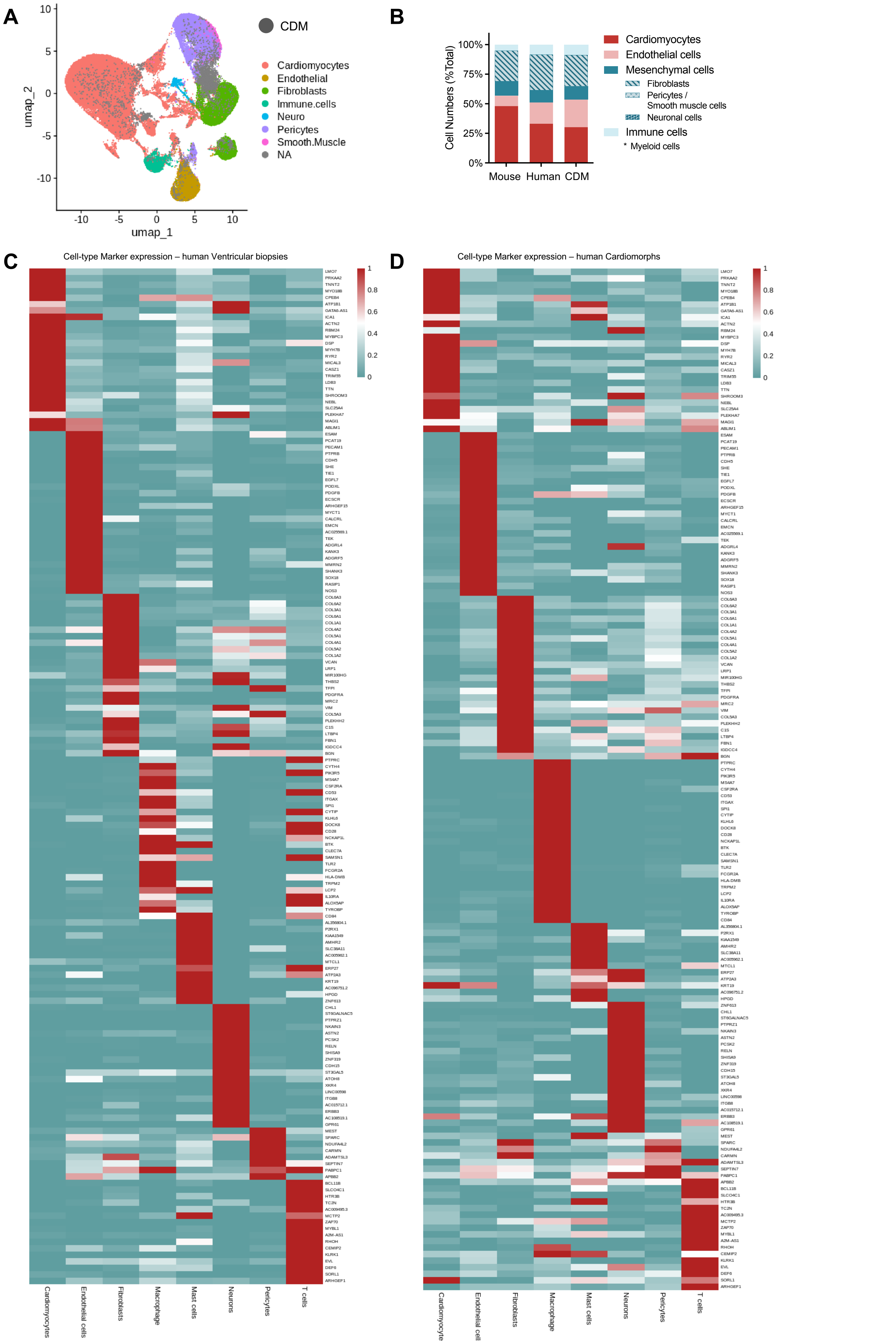

### Extended Data Fig. 4

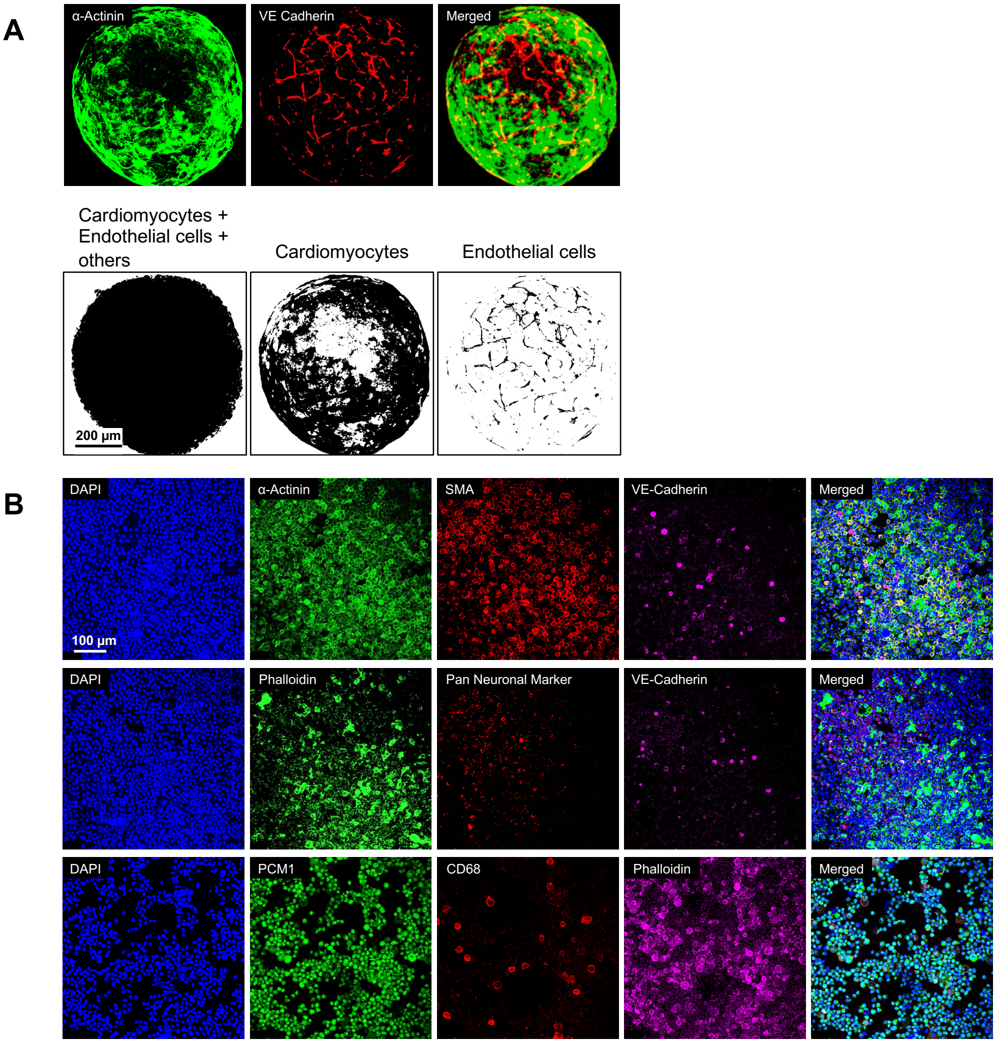

#### Extended Data Fig. 5

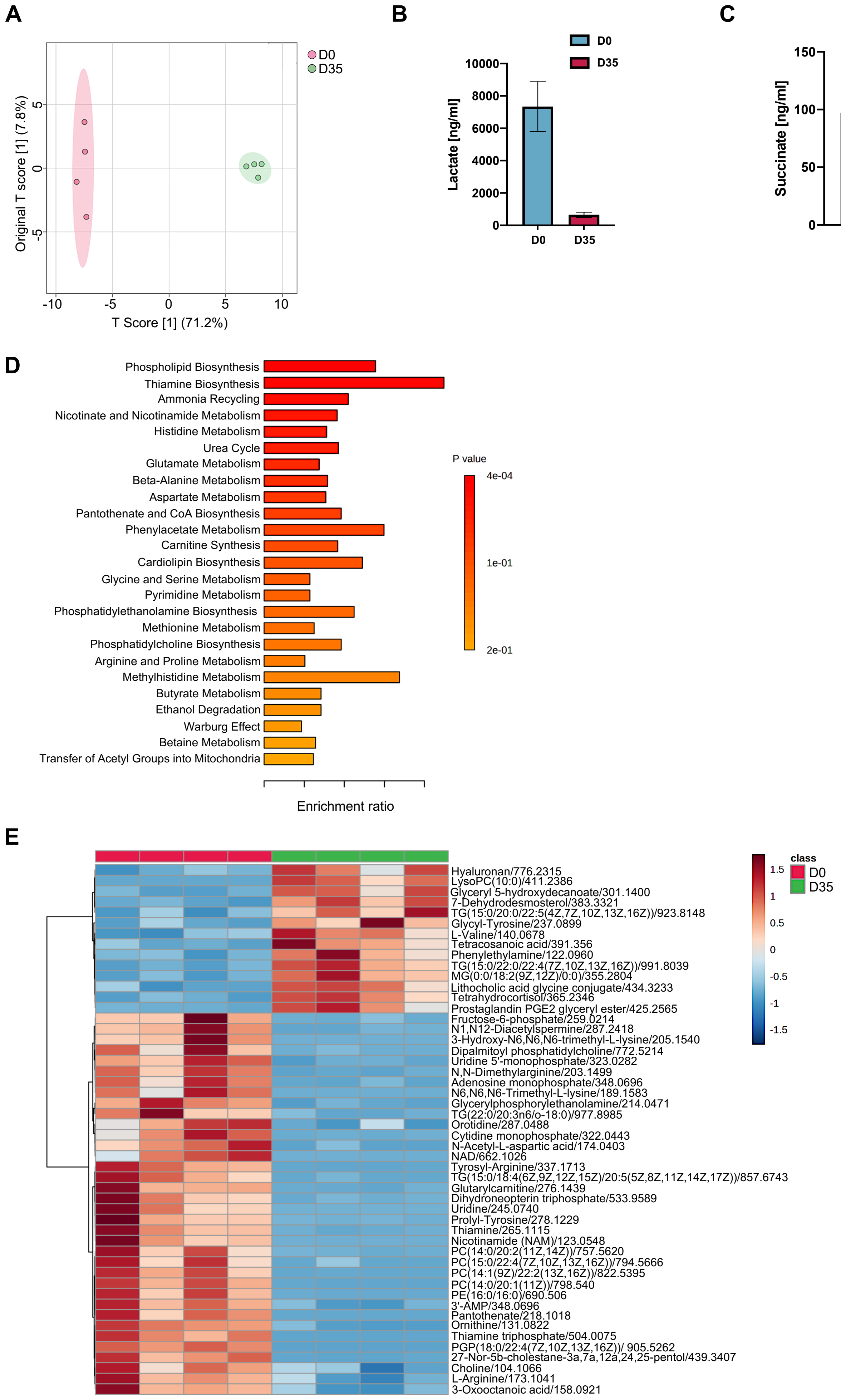

### Extended Data Fig. 6

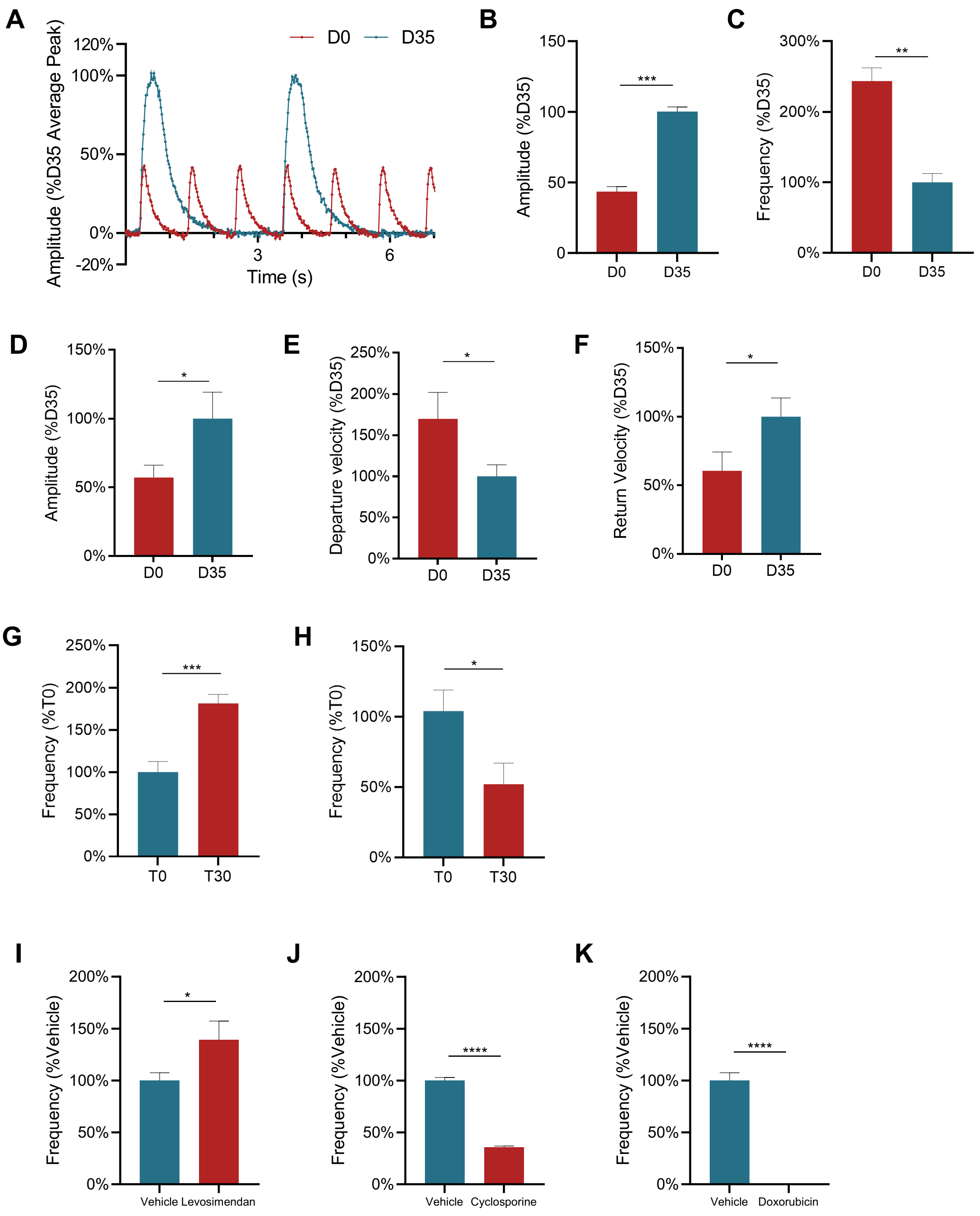

### Extended Data Fig. 7

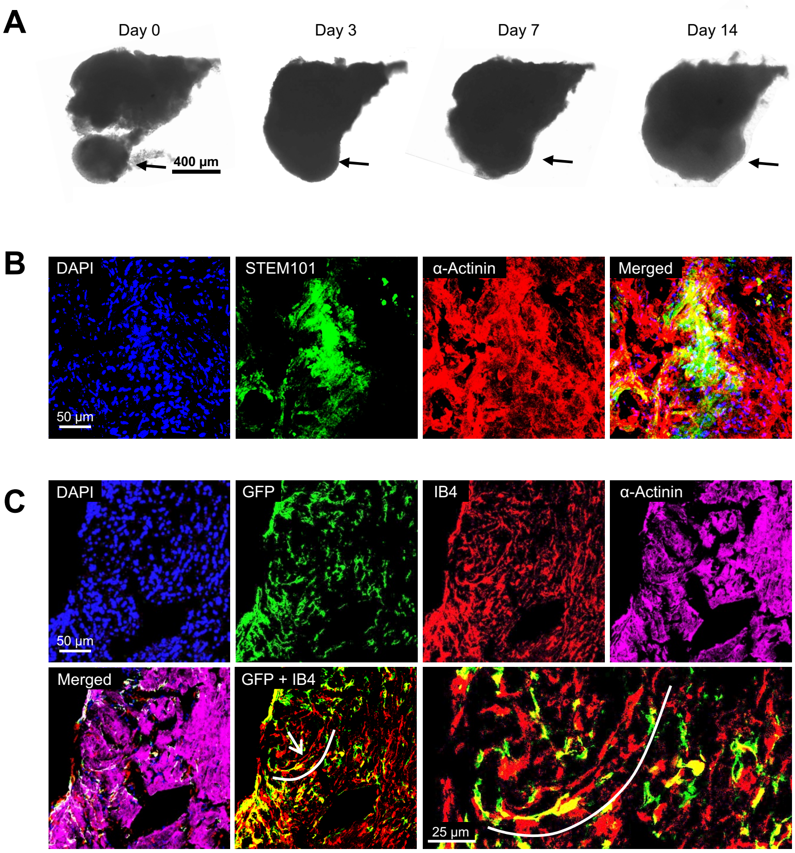

Extended Data Fig. 8

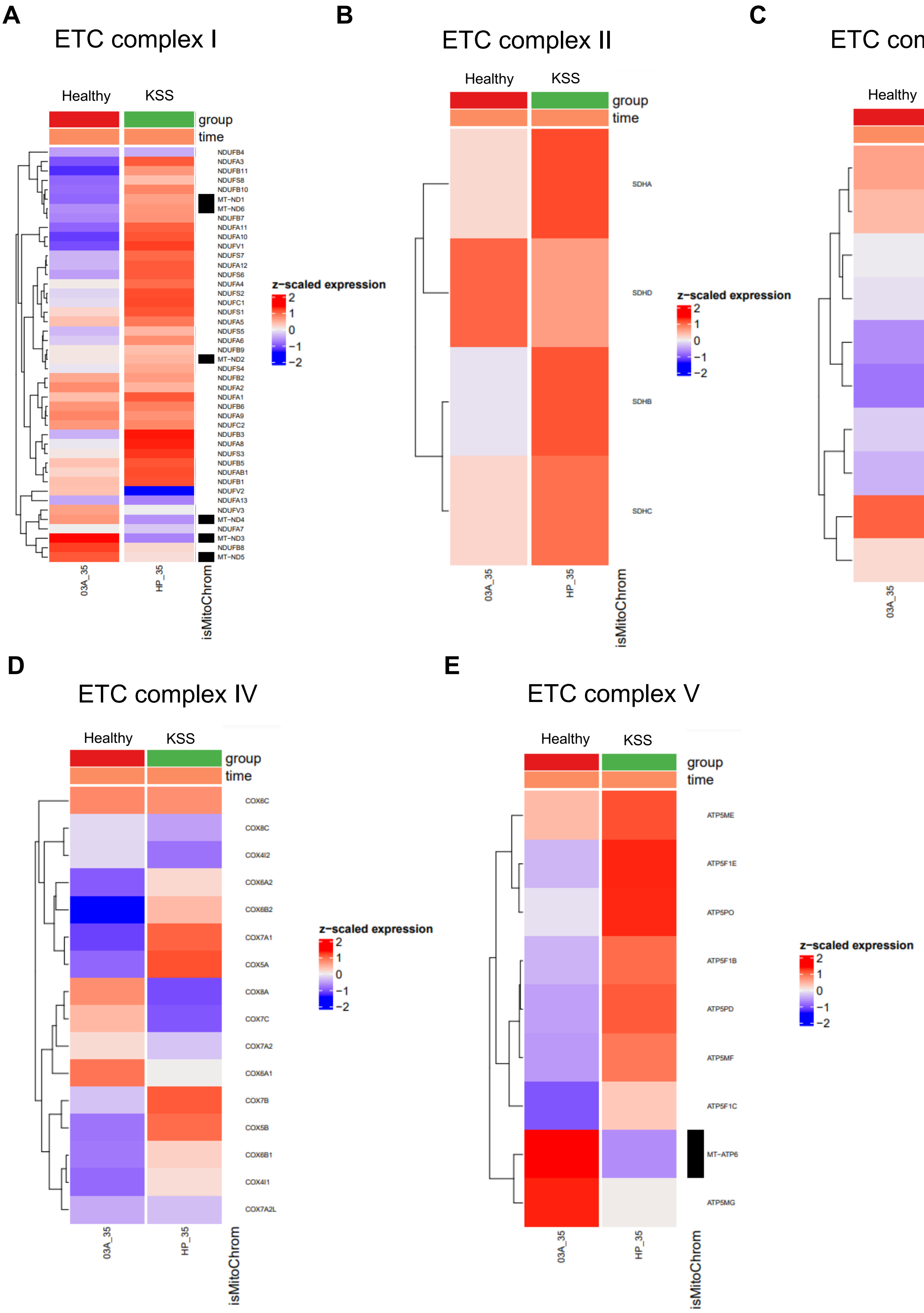

### Extended Data Fig. 9

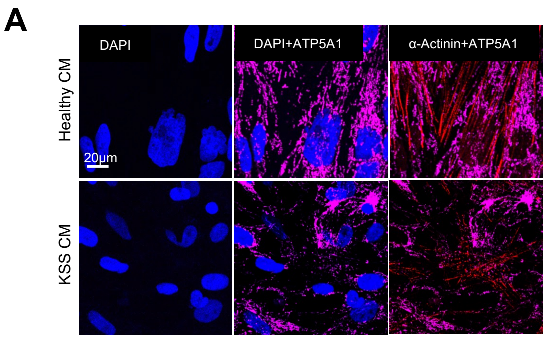

#### Extended Data Fig. 10

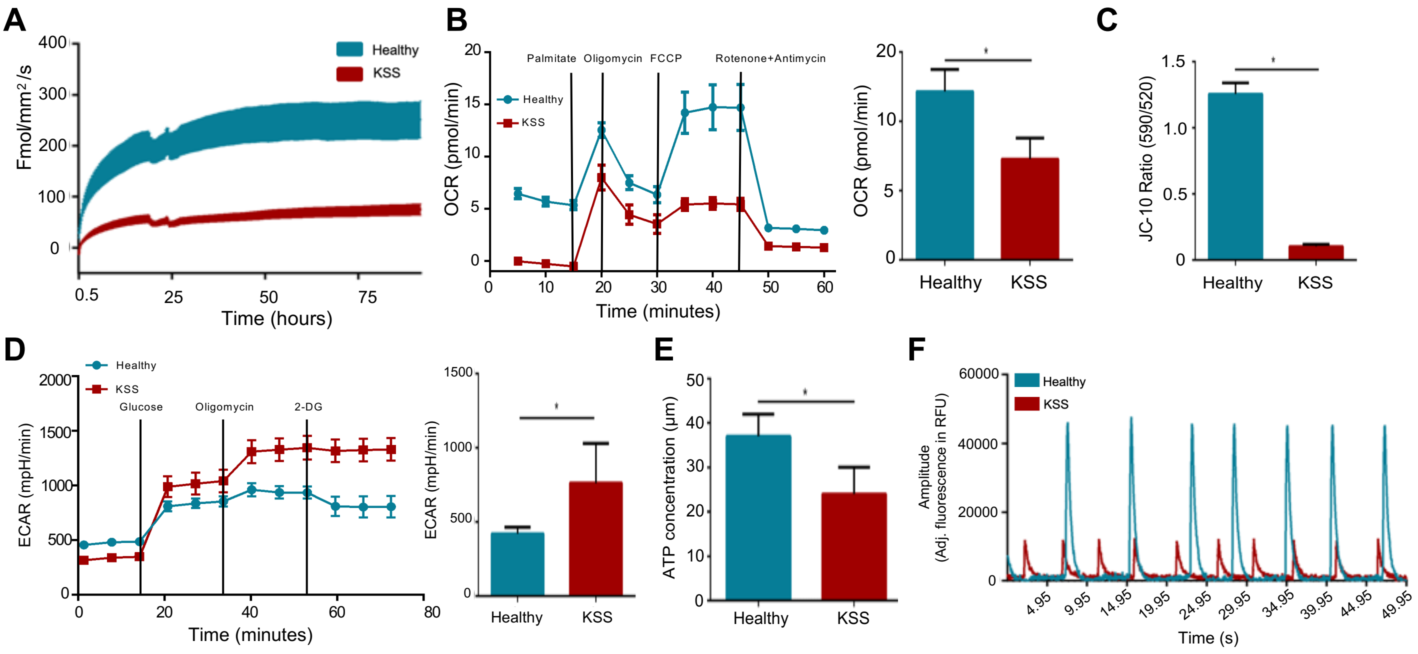

**Extended Data Fig. 11**

**A**

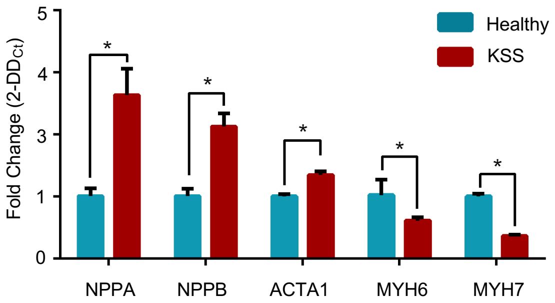


**B**

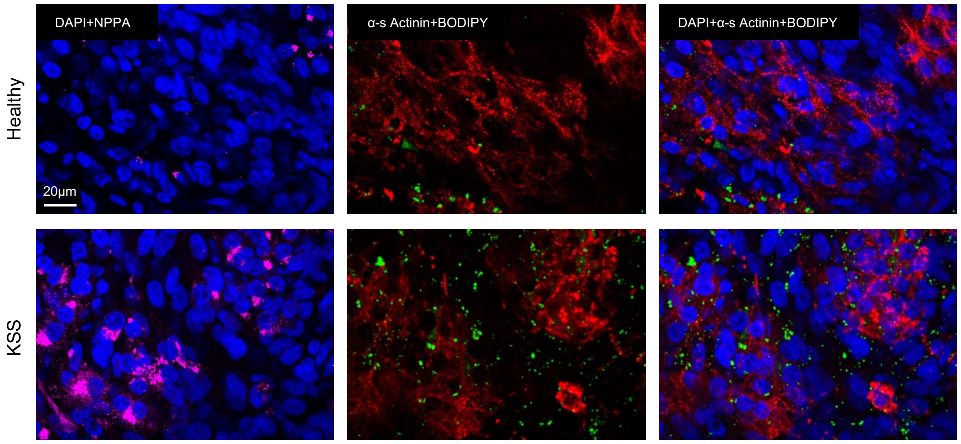

### Extended Data Fig. 12

**A**

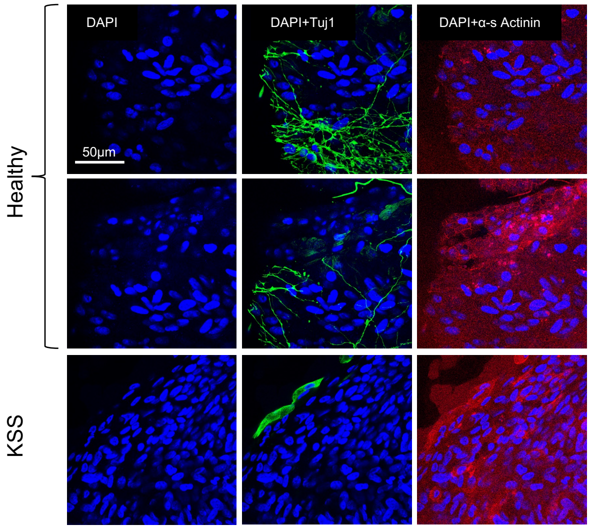


**B**

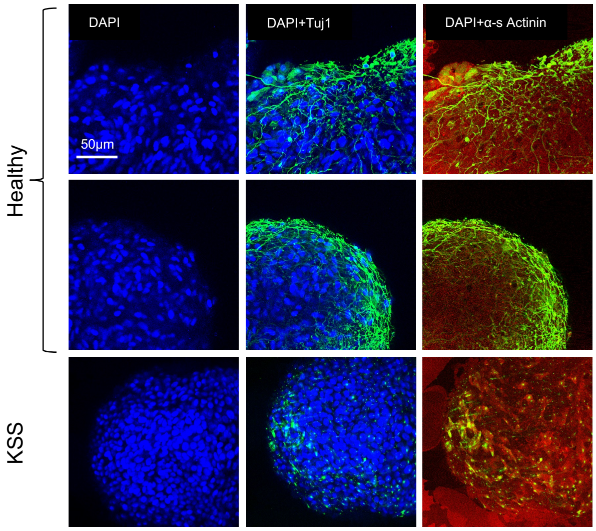

**Extended Data Fig. 13**

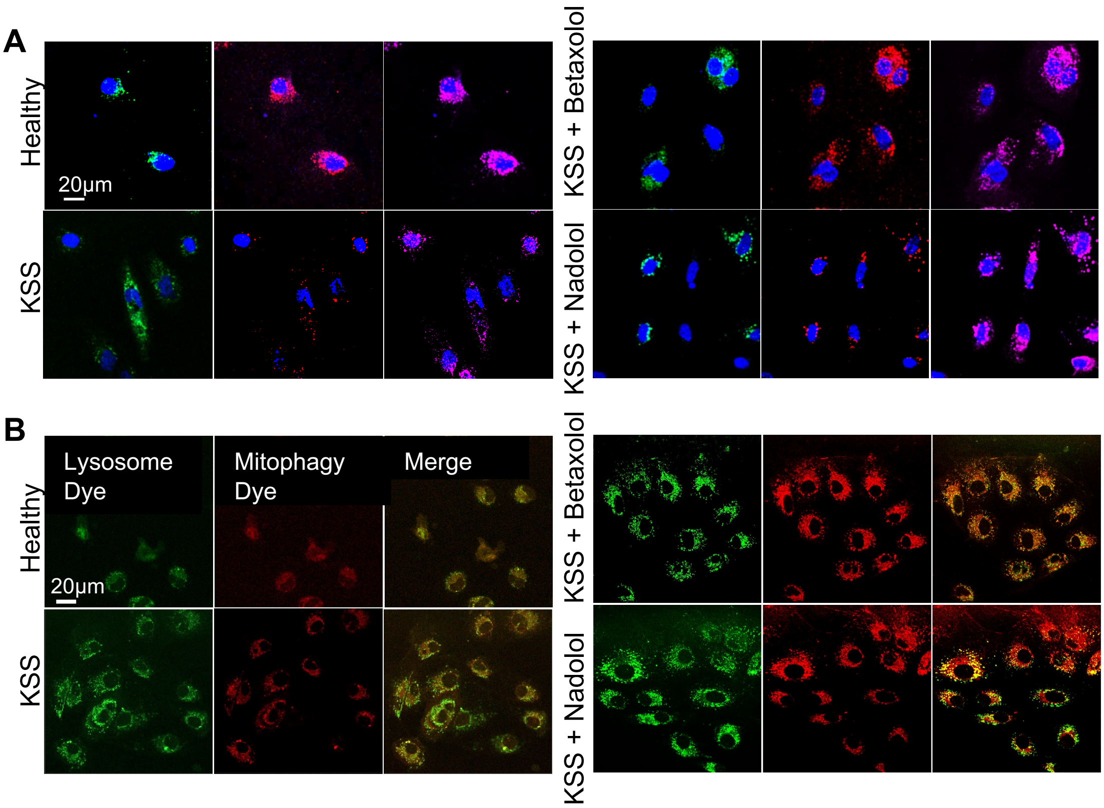
